# Chiral Graphene Quantum Dots Enhanced Drug Loading into Exosomes

**DOI:** 10.1101/2023.01.20.523510

**Authors:** Youwen Zhang, Yini Zhu, Gaeun Kim, Ceming Wang, Runyao Zhu, Xin Lu, Hsueh-Chia Chang, Yichun Wang

## Abstract

As nanoscale extracellular vesicles secreted by cells, exosomes have enormous potential as safe and effective vehicles to deliver drugs into lesion locations. Despite promising advances with exosome-based drug delivery systems, there are still challenges to drug loading into exosome, which hinder the clinical applications of exosomes. Herein, we report an exogenous drug-agnostic chiral graphene quantum dots (GQDs) exosome-loading platform, based on chirality matching with the exosome lipid bilayer. Both hydrophobic and hydrophilic chemical and biological drugs can be functionalized or adsorbed onto GQDs by π–π stacking and van der Waals interactions. By tuning the ligands and GQD size to optimize its chirality, we demonstrate drug loading efficiency of 66.3% and 64.1% for Doxorubicin and siRNA, which is significantly higher than other reported exosome loading techniques.

## INTRODUCTION

Exosomes have been extensively explored as drug delivery vehicles due to their biological and functional characteristics such as low immunogenicity, long circulation time, nontoxicity, optimal biocompatibility, strong tissue penetration, enhanced targeting effect, and ability to cross the blood-brain barrier.^1^ However, production and clinical applications of exosome-based drug delivery vehicles remain elusive due to drug loading challenges. Exosome drug loading efficiencies of the available approaches are relatively low.^2,3^ Most of these approaches potentially result in lipid damage, protein denaturation, and precipitation of nucleic acids.^4^ Thus, efficient loading of various exogenous therapeutics into exosomes is gaining increasing recognition.^5^

Endogenous drug loading involves specific cell cultures or transfected/programmed cell cultures that secret drug-loaded exosomes.^2^ It does not require any auxiliary loading apparatus and is hence quite simple. However, it cannot be used for chemical drugs and the yield of secreted exosomes with biologics is generally low (< 30%)^6^ due to the myriad of intracellular exosome biogenesis and molecular sorting mechanisms. Active exogenous drug loading methods such as sonication, electroporation, extrusion, surfactant (saponin) permeabilization, liposome fusion and freeze-thaw cycles aim to incorporate the drugs after exosome harvesting and isolation.^7^ As these active methods involve disruption of the exosome bilayer, they can also lead to exosome lysis or fusion. Hence, the size distribution, function, zeta potential and drug capacity of the cargo-loaded exosomes are sensitive to the loading procedures.^8^ The most popular electroporation and sonication methods^9^ can also damage the loaded cargos, leading to lipid degradation, protein denaturation and nucleic acid precipitation due to acoustophoretic and electrophoretic effects and their related dipolar force fields at the single-molecule level.^10, 11^

Passive exogenous loading by incubation of drugs and extracted exosomes^12^ is the least disruptive loading technology among the currently available methods. It is also the most scalable to large-volume production relevant to clinical translations and applications.^13^ However, with just diffusion across the lipid bilayer as the only loading mechanism, passive incubation can only be applied to soluble drugs with lipophilic moieties.^14^ Consequently, small interfering RNAs (siRNA) cargoes are often conjugated with hydrophobic molecules to facilitate this transport. However, other than their potential toxic effects, such moieties often prevent luminal drug loading with the drug either adhering to the exosome or intercalated at its bilayer.^14, 15^ Without the conjugation of hydrophobic moieties, passive loading efficiency of soluble drugs is typically lower than 10%.^16, 17^

Clearly, an easily optimized high-efficiency passive drug loading carrier that can adhere to the exosome bilayer but also permeate into the exosome luminal interior would be ideal. Due to their favorable hydrophobic and van der Waals interactions with lipid bilayer, graphene nanosheets has been suggested as a viable drug carrier that allows intercalation with the exosome lipid bilayer.^18^ The delocalized electron of the graphene allows easy adsorption of hydrophobic drugs and hydrophilic drugs can be functionalized directly onto the graphene. However, graphene drug carriers are often encapsulated within the bilayer without transit into the interior. In a recent study^19, 20^, we discovered a potential mechanism that allows this final transit into the interior of the exosomes–chiral interaction between the graphene sheets and the lipid molecules.

As chirality is hard-wired into every biological system,^21^ chirality of nanoparticles (NPs) play a key role in their biomedical applications involving biological events, such as cell uptake, immune response, and tissue transport.^22^ It was demonstrated that NPs coordinated with *D-*chirality displayed threefold higher permeability through cell membrane penetration in breast, cervical, and multiple myeloma cancer cells than those with *L*-chirality.^23^ These *D*-formed NPs exhibited superior stability and longer biological half-lives *in vivo*.^24^ Graphene quantum dots (GQDs), as single-layered graphene nano-sheets, have widely been used in biomedical applications due to its low cytotoxicity and high biocompatibility,^25^ unique optical properties,^26^ precisely controlled physical and chemical properties.^27^ Most importantly, GQDs displayed superior passive transport properties^28^ through cellular lipid membrane due to their two dimensional (2D) morphology,^18^ chemical structure,^29^ and nanoscale dimensions.^25^ In particular, GQDs with right-handed chirality (*D*-GQDs) derived from *D*-cystine^19^ was previously reported to permeate phosphatidylcholine (POPC) lipid bilayers more efficiently compared to the ones with left-handed chirality (*L*-GQDs).^20^ In addition, GQDs are capable of carrying a wide range of drugs with a high efficiency (>90%), and these drugs carried by GQDs with arbitrary charge can be hydrophobic or hydrophilic, biological or chemical via pi stacking and van der Waals interactions.^30, 31^ Therefore, chiral GQD with right handiness is a potential tool to efficiently load drugs/genes into exosomes by passive transport through the exosomal lipid biolayers that is similar to the membranes of the parent cells.^32,33^

In this work, we investigated permeability of chiral GQDs into exosomes through lipid membrane and their ability of passive drug loading in exosomes. (**Figure 1a**) We found that *D*-GQDs derived from *D*-cysteine have a stronger tendency (52.4 ± 8.2 % of permeation efficiency) to permeate into exosomes than pristine GQDs and *L-*GQDs, and the permeability of *D*-GQDs could further be improved to 85% after optimization. Taking advantage of chiral interactions between chiral NPs and biological lipid membranes, and the synergistic effect of physical surface of chiral GQDs, we successfully loaded a hydrophobic chemotherapy drug -Doxorubicin (Dox) and a large biological drug -siRNAs into exosomes. Dox/siRNA loaded exosomes exhibited effective killing of cancer cells, with significant knock-down of the targeted gene and the inhibited expression of relative protein levels, respectively. Our work indicated that passive drug loading into exosomes enhanced by chiral GQDs is hence the most robust (drug-agnostic) and scalable loading method that requires minimal tuning. Therefore, our developed chiral GQD enhanced drug loading technology provides a promising platform for loading therapeutic exosomes for future clinical translation.

**Figure 1.**
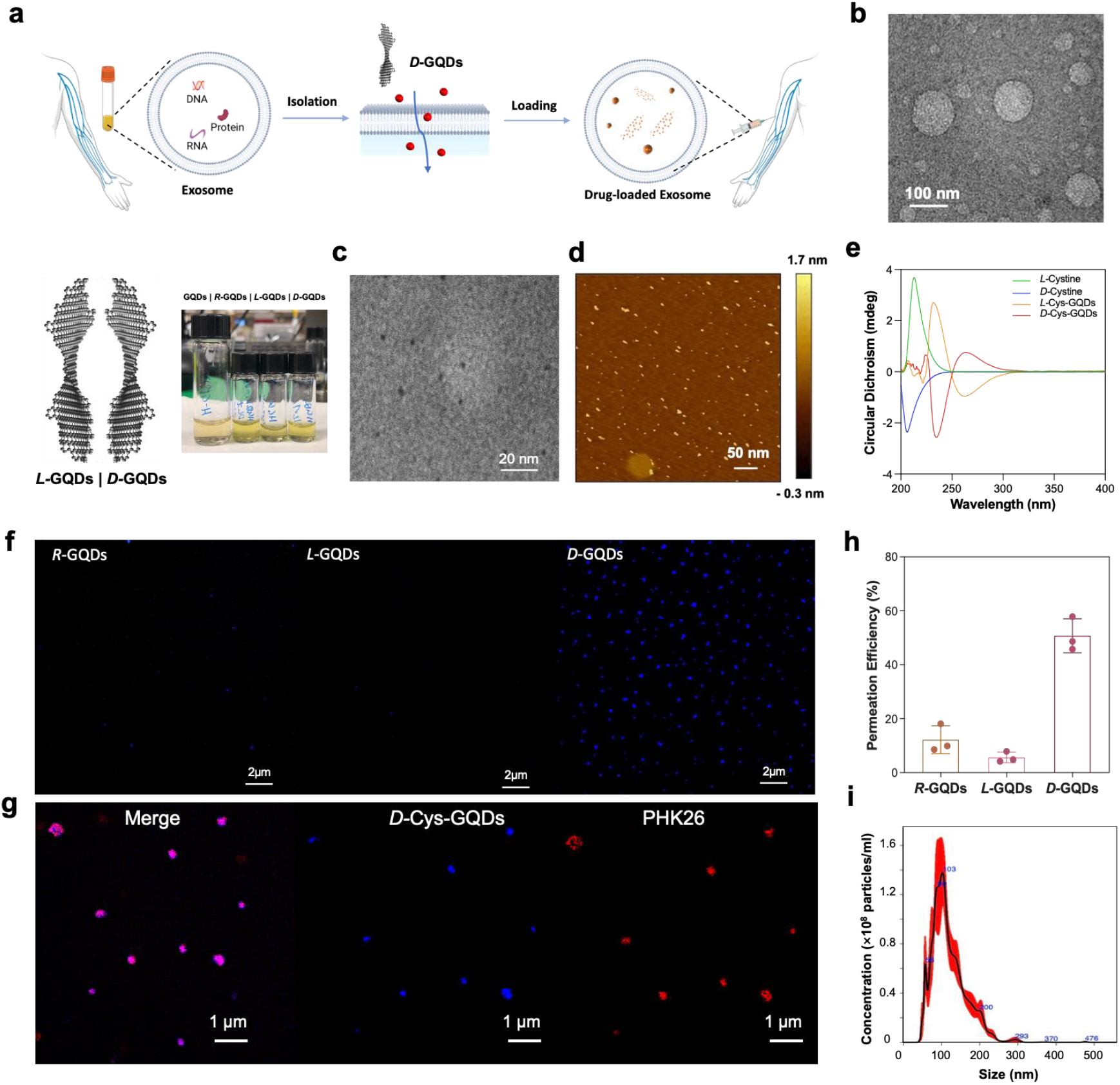
(a) The principle of chiral graphene quantum dots (GQDs enhanced drug loading into exosomes. (b) Transmission electron microscope (TEM) images of isolated exosomes by asymmetric nanopore membrane (ANM). (c) Characterization of Chiral GQDs by transmission electron microscope (TEM), (d) Atomic force microscopy (AFM) and (e) circular dichroism (CD). (f) Permeation of chiral GQDs (Blue) into exosomes was evaluated by confocal microscope. (g) Confocal images of PHK26 labelled exosomes (Red) were taken after treated with *D-*Cys-GQDs (Blue). (h) Permeation efficiency was quantified by counts of GQDs loaded exosomes (Blue) over the total counts of exosomes. (i) Size distribution and particle number of GQD-loaded exosomes were measured with nanoparticle tracking analysis (NTA).

## Results and Discussion

### Investigation of Chiral GQD Permeation into Exosome

To investigate the permeation efficiency of chiral GQDs into exosomes (**Figure 1a, b**), we first synthesized chiral GQDs derived by surface modification with *L/D*-cysteine(*L/D*-Cys-GQDs) using a previously reported method.^19^ The structures of derived *L/D*-Cys-GQDs were confirmed by the combination of spectroscopy and microscope. Transmission electron microscope (TEM) images showed that *L/D*-Cys-GQDs has a size distribution of 3.0 - 9.4 nm (**Figure 1c** and **Figure S1a**). Atomic force microscopy (AFM) images (**Figure 1d**) confirmed the size range of *L/D*-Cys-GQDs in TEM. The thickness of *L/D*-Cys-GQDs was within 2 nm verified by AFM (**Figure 1d**), which was corresponding to a single layer of chiral GQDs with enhanced height from helical buckling (twisting) of the pristine GQDs (∼1 nm).^34, 35^ *L*- and *D*-Cys-GQDs showed positive and negative peaks at 236 nm respectively (**Figure 1e**) in circular dichroism (CD) spectra. These chiroptical bands and *g*-factor (**Figure S1b**) of chiral GQDs showed opposite signs depending on the reveal of right- and left-handed twists of the 2D nanosheets respectively, matching the chirality of *L/D*-cysteine used for conjugation to the GQDs.^19^ Cloud peaks at 264 nm indicated covalent bonding of cysteines on the edge of GQDs.^19^ Whereas the as-synthesized GQDs, and *R*-Cys-GQDs (i.e., made with racemic mixture of *L*- and *D*-cysteine) displayed no chiroptical activity in CD spectra (**Figure S1c**) at either a high-energy peak at 220−250 nm or a low-energy peak at 250−300 nm. Furthermore, the UV-Vis spectra (**Figure S1d**) of the chiral GQDs revealed two distinct absorption bands centered at 270 nm (ascribed to *π– π\** transitions in the *sp*^2^-hybridized carbon core) and 375 nm (attributed to the *n–π*\* transitions or carboxyl groups). It indicated partial relaxation of exciton confinement compared to pristine GQDs with a peak at 225 nm due to the hybridization of the aromatic system of GQDs with the atomic orbitals on cysteine moieties.^19^ Being excited by photons with λ_ex_ = 360 nm, pristine GQDs showed the emission at 480-540 nm, while *L*-, and *D*-forms of GQDs displayed strong emission at 500−550 nm (**Figure S1e**). The red shift (∼ 26 nm) after the modification with the amino acids in the photoluminescent (PL) peaks was caused by charge transfer between functional groups and the graphene carbon core of GQDs, which narrowed the band gap. ^36, 37^. The zeta-potential (ζ) of as-synthesized GQDs was −21.8 ± 4.2 mV (**Figure S1f**). After attachment of cysteine moieties, ζ-potentials of *L-* and *D-*form chiral GQDs became −3.3 ± 2.1 mV and −1.6 ± 1.8 mV (**Figure S1f**), respectively. Such reduction of surface charge from GQD to *L/D*-Cys-GQDs was consistent with amidation of negatively charged carboxyl groups (−COOH) at the edges of GQDs while retaining a substantial degree of ionization. Taken together, the combined results supported our successful synthesis of chiral GQDs.

Following synthesis and characterization of *L/D*-Cys-GQDs, we investigated the permeation of chiral Cys-GQDs into exosomes isolated from cell culture media of 3T3 cell lines (**Figure 1b** and **Figure S2**). As-synthesized GQDs and their chiral derivates (7.5 μM) were incubated with 3T3 exosomes (1.09×10^9^ particles/mL) in phosphate-buffered saline (PBS) buffer (pH 7.4) at room temperature under a static condition. The permeation of GQDs and their chiral derivatives into exosomes were examined by confocal imaging based on the innate fluorescence of GQDs at 480 - 540 nm. Exosomes after incubation with GQDs were washed with PBS buffer under the hold of 100 kDa centrifuge tube until the fluorescence intensity of GQDs in the mixture decreased to a constant (**Figure S3**). The permeation of chiral GQDs into exosomes was detected by the accumulation of innate fluorescence of GQDs (around 525 nm, marked as blue) in the exosomes using confocal microscope. Cys-GQDs with right-handed chirality (*D*-Cys*-*GQDs) showed the significantly higher density of blue dots (the accumulation of GQDs) than achiral *R-*Cys*-*GQDs and left-handed *L-*Cys*-*GQDs (**Figure 1f**) under the same concentration of the exosomes (10^8^ particles/mL). To verify that the observed fluorescent signal of GQDs were inside exosomes, we labelled exosomes by staining their membranes using PKH26 that is a red fluorescent lipid membrane dye. The colocalization of PHK26 labeled exosome (Red) and *D-*Cys*-*GQDs (Blue) in **Figure 1g** demonstrated the permeation and accumulation of *D*-Cys*-*GQDs in the exosomes. To note, some point source (100 ∼ 500 nm) in the fluorescent image was about twice larger than the actual size of the objects (30 ∼ 200 nm) due to the diffraction effects and resolution limitation of optical microscope (180 nm laterally and 500 nm axially),^38^ thus they were within the dimension range of exosomes. In order to quantify the permeation efficiency of GQDs into exosomes, we developed a counting method based on image analysis (**See Method**) to reflect the loaded exosomes at single exosome level. In short, the permeation efficiency of GQDs in exosomes were determined by statistically analyzing the total amount of GQDs loaded exosomes (blue dots larger than 30 nm as a threshold) over the total counts of exosomes from nanoparticle tracking analysis (NTA). *D*-Cys-GQDs exhibited significantly higher permeating efficiency (52.4 ± 8.2%) than *R*-Cys-GQDs (13.8 ± 4.7%), while *L-*Cys*-*GQDs (5.7 ± 2.2%) showed very limited permeation (**Figure 1h**). Most importantly, the size and morphology of exosomes retained integrity after the loading procedure by *D-*Cys-GQDs based on TEM and NTA results (**Figure S4** and **Figure 1i**). This result aligned with the previous findings by Molecular Dynamics (MD) simulation that *D*-Cys-GQDs had a stronger tendency to penetrate the cellular lipid membrane of mammalian cells, than *L-*Cys*-*GQDs.^19^ Such transport phenomenon is potentially attributed to the twist of 2D nanosheet that gives rise to nanoscale chirality with single chiral center,^39^ resulting in the interactions between chiral GQDs and lipid membrane of exosomes.

### The effect of size on permeability of chiral GQDs into exosomes

The interaction of GQDs with lipid bilayer membrane depends on the size of GQDs.^20^ In particular, small GQDs (< 13.3 nm) are able to enter the lipid bilayers while maintain the membrane intact.^20, 40^ However, larger GQDs tend to deform the membrane with the formation of hemisphere vesicles and cause potential damage.^41, 42^ Moreover, the chirality originated by lattice distortion at nanoscale are influenced by the size of NP lattices,^43^ thus may affect the permeation efficiency of chiral GQDs in exosomes. Here, we investigated the effect of *L/D*-Cys-GQDs with three different sizes on their permeation into exosomes. The average sizes of the three GQD samples were 5.14 nm, 25.6 nm and 65.7 nm (**Figure 2a, b**) in this study. Due to the same tendency of fluorescence spectra and CD spectra (**Figure S5a**) of *L/D*-Cys-GQDs, *D-*Cys-GQDs that has higher permeation efficiency was demonstrated in **Figure 2** to show the size effect. With the size increase of chiral GQDs, bathochromic shift was observed in fluorescent emission spectra of *D*-Cys-GQDs (exited at 360 nm, **Figure 2c**). The chirality corresponding to the nanoscale distortion was observed at a low-energy peak (250−300 nm) in CD spectra. Smaller *D-*Cys-GQDs (5.14 nm) exhibited higher chiroptical activities at 250-300 nm than the larger ones (25.6 nm and 65.7 nm). This was further confirmed by the CD peak at 265 nm of smaller *D*-Cys*-*GQDs associated with Cotton effect (**Figure 2d**).^44^ This phenomenon was potentially due to larger dihedral angles in the twisted molecular structures of the smaller chiral GQDs according to the previously reported study.^19^ In addition, there were more terminal -COOH groups on larger GQDs, which increased the probability of anomalous and asymmetric bindings of cysteines, diminishing the symmetry-breaking perturbation of chiral edge ligands to electronic states of graphene nanostructures.^19, 45^ Following the characterizations of chiral Cys-GQDs with different sizes, we investigated the permeation of these *L/D*-Cys-GQDs into exosomes isolated from 3T3 cell culture media by ultracentrifugation method. The largest chiral GQDs (65.7 nm) showed limited permeation into exosomes according to quantification method of permeation efficiency in the previous session (**Figure 2e**). This was potentially because the size of the largest GQDs (65.7 nm) was comparable to the size of exosomes (116 nm, **Figure S1**). Instead of passive transport through the membrane, the large GQDs damaged the lipid membranes of exosomes, confirmed by TEM images of exosomes (debris of exosome observed in **Figure S5b**). Compared to the smaller *D-*Cys*-*GQDs (5.14 nm) with a permeation efficiency of 82.7% into exosomes, the ones with a median size (25.6 nm), has a low permeation efficiency (16.9 %). This is consistent with the size distribution of the median GQDs, of which, 9.4 % are below 13.3 nm. The permeation efficiency of *D-*Cys*-*GQDs was 1.5- to 3-time higher than that of *L-*Cys*-*GQDs for both sizes of 5.14 nm and 25.6 nm, indicating the chirality-selective passive diffusion of GQDs through exosomal membranes. Overall, the results suggested that small *D-*Cys*-*GQDs can permeate 3T3 exosome membranes most efficiently, which can be attributed to the relative size of GQD to exosomal membrane thickness^41, 42^ and size-associated chirality generated from distortion of NP lattices. ^43, 46, 47^

**Figure 2.**
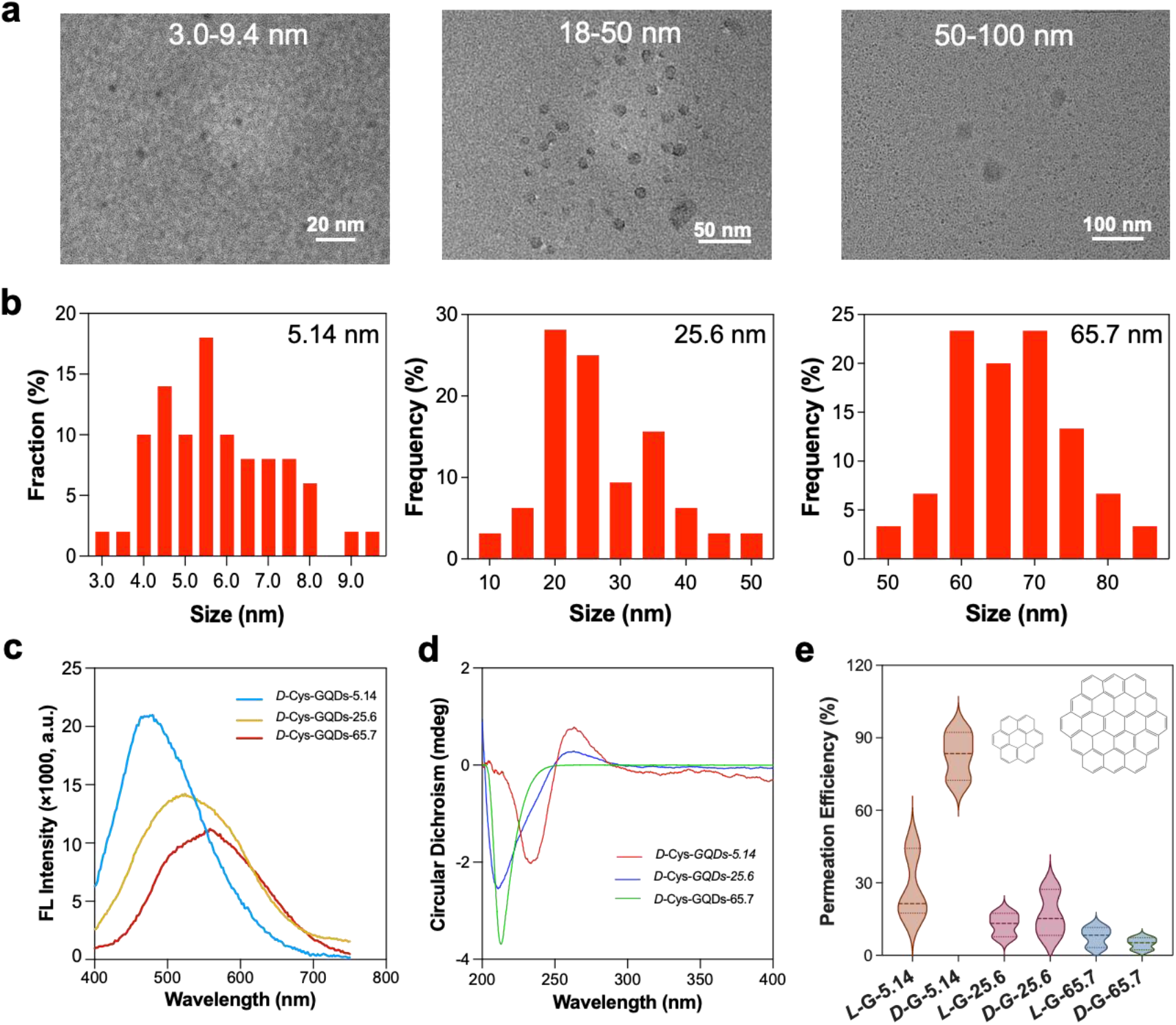
Size effect on permeation of chiral GQDs into exosome. (a) TEM images and (b) histograms of size distribution of three size ranges of GQDs. Characterization of chiral GQD under different size ranges, (c) fluorescent spectra exited at 360 nm, (d) CD spectra, and (e) permeation efficiency of size-dependent chiral-GQDs (15 μM) into exosome (*L*-G: *L-*Cys-GQDs and *D*-G: *D-*Cys-GQDs).

### The effect of ligand on permeability of chiral GQDs into exosomes

To investigate the ligand effect on permeation efficiencies of chiral GQDs into exosome, GQDs were functionalized with another two chiral amino acids, arginine and tryptophan, by the same coupling reaction for *L/D-*Cys-GQDs to generate different chiral GQDs (**Figure 3a**). The successful conjugations of arginine and tryptophan onto the edge of GQDs were supported by UV–vis absorption and fluorescent spectra. *L/D-*Arg-GQDs and *L/D-*Trp-GQDs both had UV–vis absorption peaks at around 270 nm and 375 nm, similar to Cys-GQDs (**Figure S6a**). When excited at 360 nm, *L/ D-*Arg-GQDs displayed strong emission at 400−500 nm (**Figure 3b**). A blue shift (∼40 nm) was observed compared to as-synthesized GQDs in the PL peaks due to the coupling of electron-donating sidechain that played a role of chromophore on GQDs. *L/D-*Trp-GQDs showed a broad range of fluorescent emission compared to Cys-GQDs and Arg-GQDs with relatively lower intensity under the same concentration (7.5 μM). This was attributed to the fact that conjugating the large indole group of tryptophan to GQDs weakened the optical property (lower intensity of emission)^48^ and the increasing the size of GQDs changed the band gaps.^49^ The chirality of these functionalized GQDs was determined and analyzed by CD spectra. CD spectra of both *L/D-*Arg-GQDs and *L/D-*Trp-CQDs gave rise to a new symmetrical CD signal at around 240-300 nm. These new peaks were different from those of free Arginine and Tryptophan near 220 nm, indicating the successful synthesis of chiral-GQDs with chiroptical activity (**Figure 3c, d**). The zeta-potentials (ζ) of chiral Arg-GQDs were +0.5 mV(*L*), +0.3 mV(*D*) and Trp-GQDs ere −1.1 mV(*L*), -0.7 mV(*D*) (**Figure S6b**).

**Figure 3.**
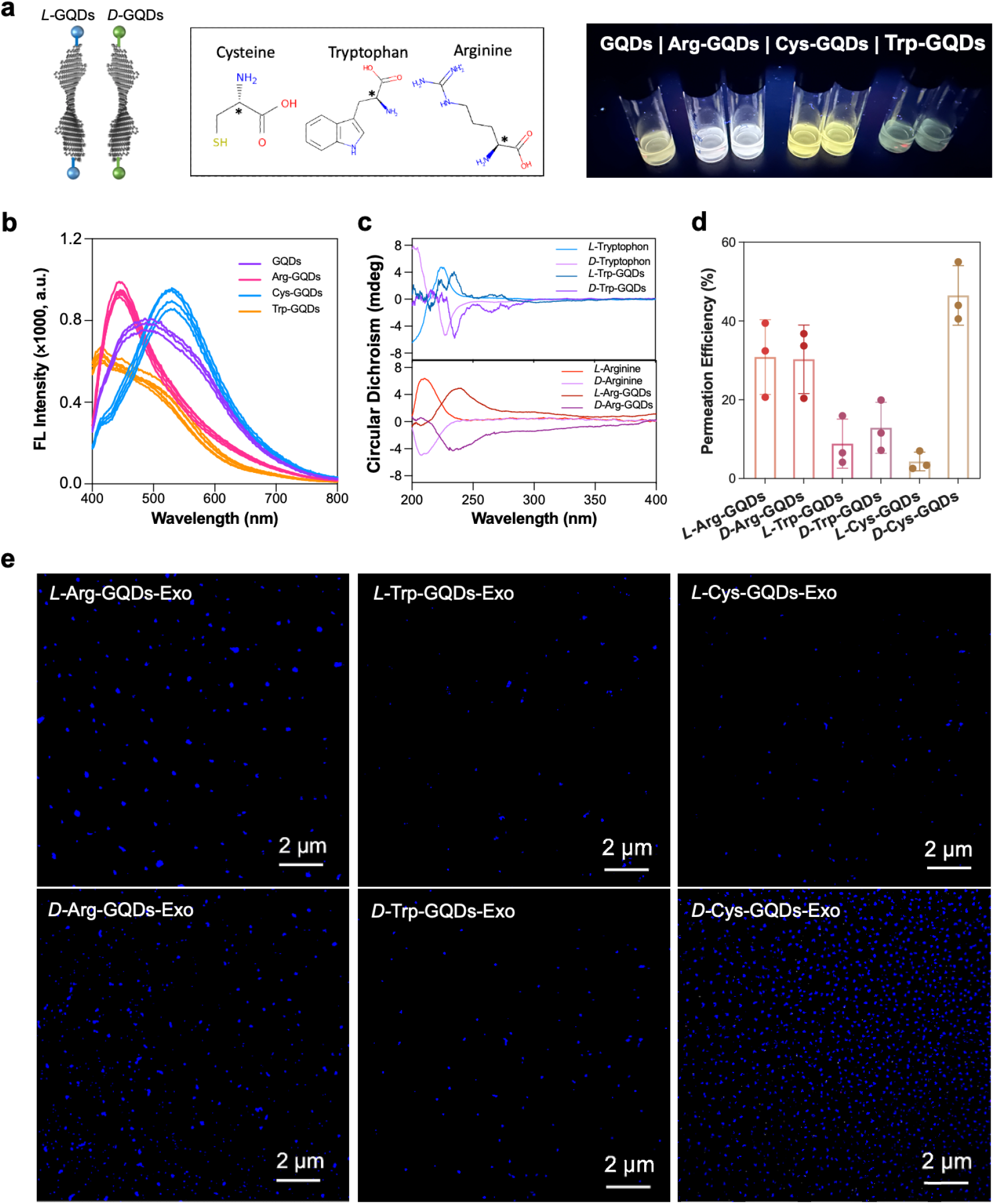
The effect of ligand on permeability of chiral GQDs into exosomes. (a) Photographs of the chiral GQDs depending on various ligands excited by UV light (λ_max_ = 365 nm). Characterization of GQDs and their chiral derivatives in (b) fluorescent spectra exited at 360 nm, (c) CD spectra, and (d, e) permeation efficiency of chiral GQDs (7.5 μM) into exosomes (10^9^ particles/mL) imaged by confocal microscope and quantified by analyzing images.

We investigated the permeation of chiral-GQDs derived from different chiral ligands into exosome under the same condition in the previous session. *D-*Trp-GQDs reflected slightly higher permeation efficiency (15.63 ± 4.39%) into exosomes than *L-*Trp-GQDs (9.79 ± 6.48 %). However, Trp-GQDs showed a significant reduction of permeation efficiency into exosomes compared with *D-*Cys-GQDs (**Figure 2d, e**). The potential reason is that the tryptophan with a large aromatic molecular structure (**Figure 3a**) gave rise to multiple chiral centers^50^ and increased the complicity of chiral interactions of chiral GQDs with lipid bilayers of exosomes. Meanwhile, Arg-GQDs presented higher permeation efficiency (*L*, 31%; *D*, 29%) (**Figure 3e**) than Trp-GQDs, however, there is no selectivity between *L-* and *D-*Arg-GQDs. From the CD spectra, the lack of Cotton effect indicated that the chirality from Arg-GQDs originated from the surface enhancement of arginine attached on GQDs non-covalently. This also reflected on the broaden bands between 275 nm and 370 nm without cloud peaks with opposite signs of ligand chirality.^51^ This is caused by the partial physical electrostatic absorption of arginine ligand on the GQDs due to the charge of amino acids (+) and as-synthesized GQDs (-). These effects further implicated that nanoscale chirality origin from twist of 2D nanosheet enhance the permeability of biological lipid membrane through selective chiral interactions. Therefore, with the highest permeation efficiency due to the nanoscale chirality,^52^ *D-*Cys-GQDs were chosen as the optimum chiral carriers for drug loading into exosomes in all the subsequent drug loading experiments.

### The effect of concentration on permeability of chiral GQDs into exosomes

Concentration gradient of NPs across the lipid membrane is one of the main driving forces for NP transport through lipid membrane.^53^ Therefore, to optimize the permeation efficiency of chiral GQDs as a drug loading vehicle, we investigated the permeation efficiency of *D-*Cys-GQDs at different concentrations in the range of 0.75–30 μM in PBS buffer. With an increase in the concentration of the *D-*Cys-GQDs, a larger amount the *D-*Cys-GQDs entered the exosomes, indicated by more accumulation of *D-*Cys-GQDs (Blue dots, **Figure 4a**) and higher retention fluorescent intensity of *D-*Cys-GQDs-loaded exosome solution (**Figure 4b**). Permeation efficiency of *D-*Cys-GQDs into exosomes enhanced with the increased concentration in the range of 0.75 to 15 μM. However, the permeation efficiency of *D-*Cys-GQDs dropped significantly when the concentration of *D-*Cys-GQDs reached up to 30 μM (**Figure 4b**). This is potentially attributed to saturation of *D-*Cys-GQDs within the exosomes that eventually resulted in damage of exosome at an external *D-*Cys-GQDs saturation concentration of 30 μM(**Figure S7**). Overall, the highest permeation efficiency, 85.2 ± 10.3 % of *D-*Cys-GQDs into exosomes was achieved at a concentration of 15 μM (**Figure 4c**). All the subsequent drug loading experiments into exosome were conducted at this optimized concentration of *D-*Cys-GQDs.

**Figure 4.**
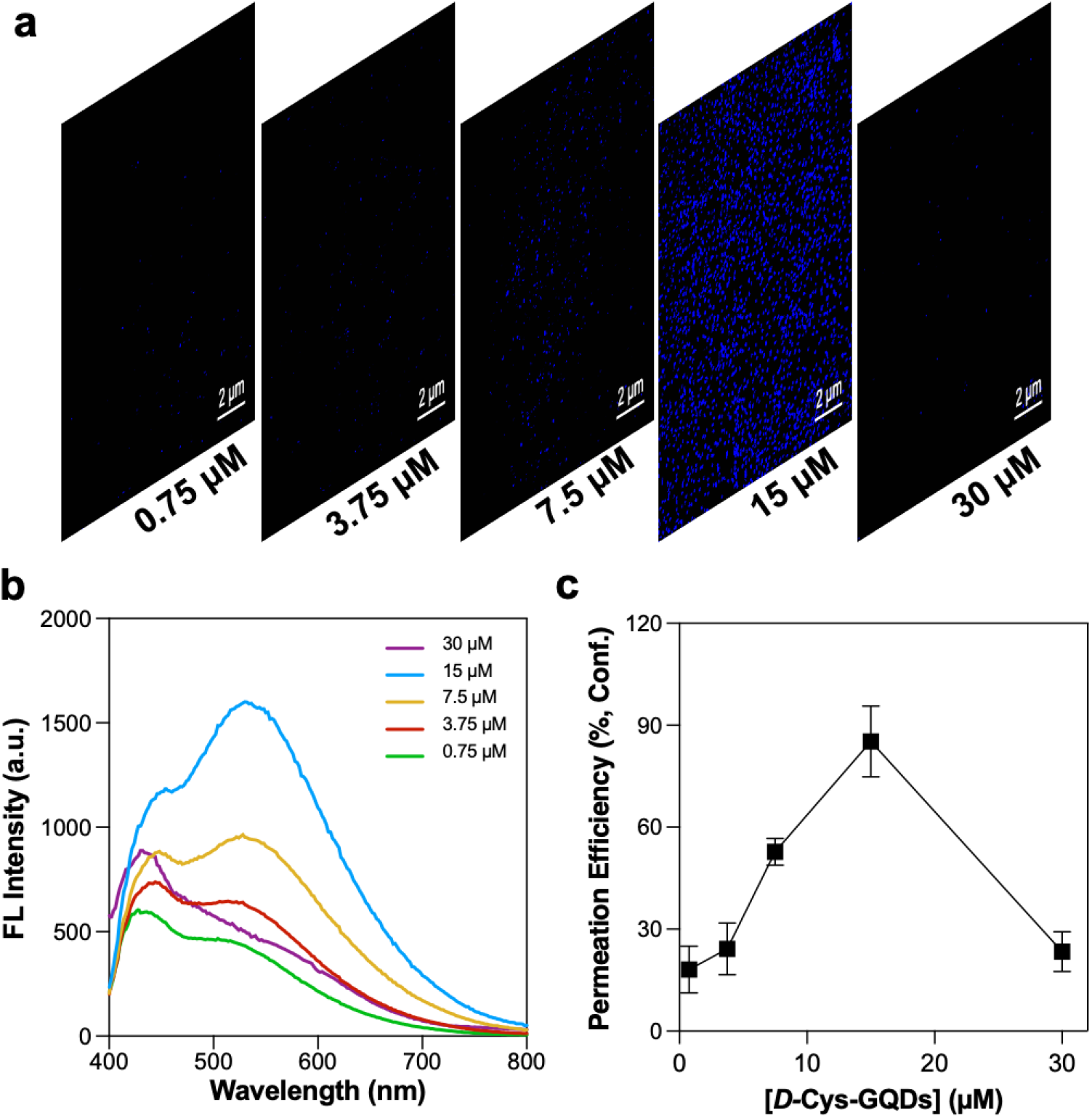
The effect of chiral GQD concentration on their permeation into exosome. (a) Confocal images of exosomes by treated with different concentration of *D-*Cys-GQDs. (b) Fluorescent spectra of *D-*Cys-GQDs loaded exosomes exited at 360 nm, and (c) statistical analysis of permeation efficiency of *D-*Cys-GQDs into exosomes.

### *D-*Cys-GQDs-enhanced chemotherapy drug loading into exosomes

After optimizing the permeability of chiral GQDs into exosomes, we evaluated the loading efficiency of a common hydrophobic chemotherapy drug, Doxorubicin (Dox), facilitated by *D*-Cys-GQDs. Dox-loaded exosomes as a promising nanomedicine have shown enhanced therapeutic effects compared to the commercially available Dox-loaded liposomes.^54^ However, loading Dox or other common drugs into exosomes remains challenging, which hinder their translational applications as drug delivery carriers.

*D-*Cys-GQDs can carry Dox molecules via π–π stacking between planar surface (carbon ring) of GQDs and anthracene group of Dox molecules. Such *D-*Cys-GQDs/Dox complex can permeate lipid bilayer of exosome membrane with high efficiency, similar to the permeation efficiency of *D-*Cys-GQDs into exosomes (**Figure 5a**). The amount of Dox molecules carried by each *D-*Cys-GQD was determined by the quenching efficiency using fluorescence resonance energy transfer (FRET) assay (**Figure 5b**).^55^ As the concentration of *D-*Cys-GQDs increased, the quenching efficiency of Dox (200 μM) decreased dramatically. The quenching efficiency of Dox began to reach a plateau when concentrations of *D-*Cys-GQDs were higher than 15 μM (**Figure S8a**), which confirmed the attachment of Dox on the surface of *D-*Cys-GQDs. Based on the molar concentrations of Dox and *D-*Cys-GQDs at the plateau of quenching efficiency, each *D-*Cys-GQD was able to carry 14 Dox molecules (**Figure S8b**). *D-*Cys-GQDs/Dox complex showed absorbance at both featured regions of *D-*Cys-GQDs and Dox, which further verify the attachment of Dox on the surface of *D-*Cys-GQDs (**Figure S8c**). Taken together, the combined results supported the formation of *D-*Cys-GQDs/Dox complex.

**Figure 5.**
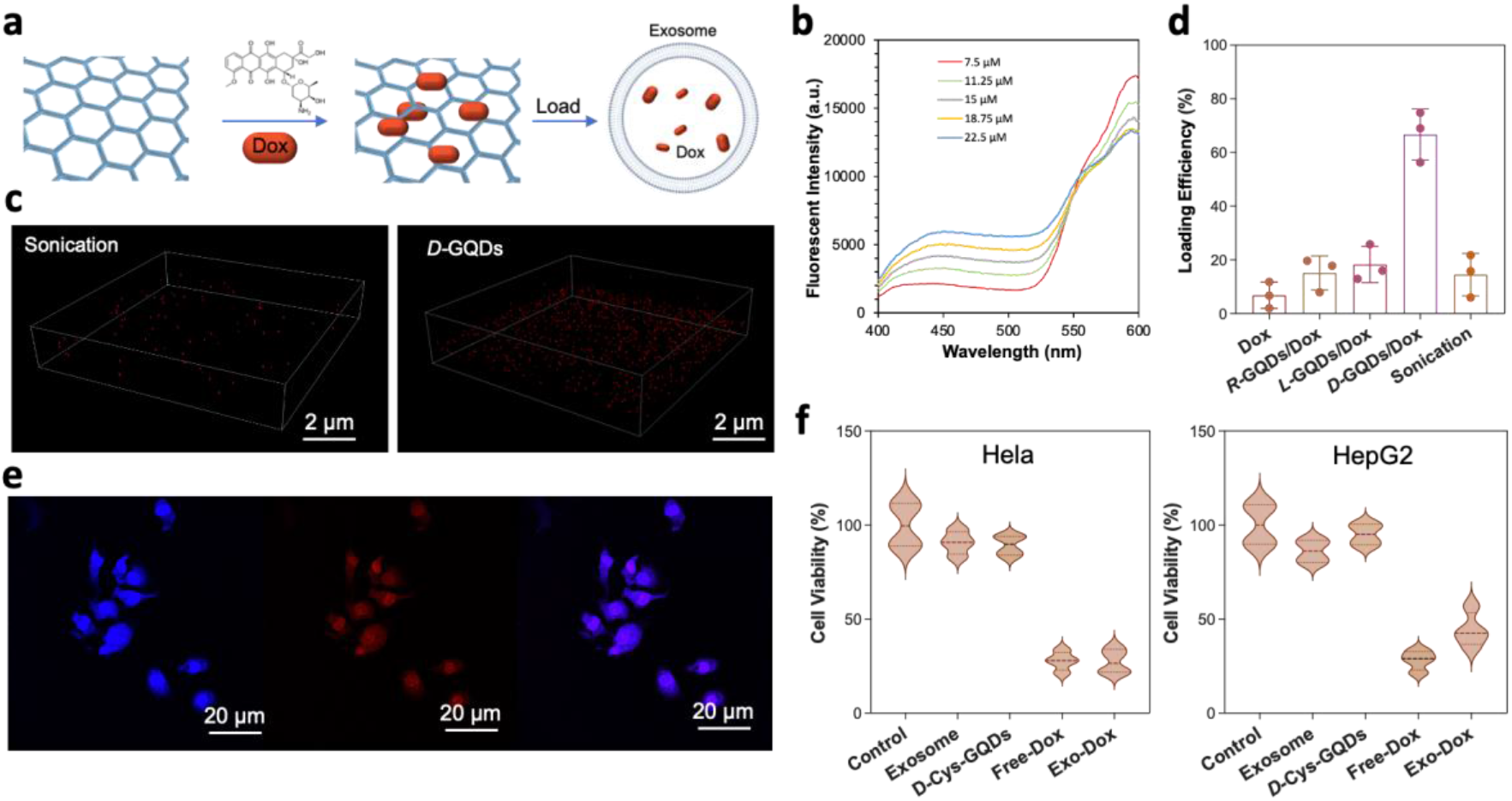
Chemotherapy drug Doxorubicin (Dox) was loaded into exosomes by *D-*Cys-GQDs. (a) Schematical illustration of Dox loading into exosomes facilitated by *D-*Cys-GQDs. (b) Characterization of attachment of Dox (200 μM) directly onto the *D-*Cys-GQDs (7.5 ∼22.5 μM) by fluorescent spectra (λ_max_ = 360 nm) and quenching efficiency. (c) Comparison of *D-*Cys-GQDs and sonication loading strategies by confocal microscope. (d) Loading efficiency of Dox into exosomes facilitated by chiral GQDs. (e) Confocal images of Exo-Dox uptake by 3T3 cells *in vitro* (Scale bars: 20 μm) and (f) cell viability was assessed using CCK-8 assay.

We investigated the loading efficiency of *D-*Cys-GQDs/Dox complex into exosomes at the optimized *D-*Cys-GQD concentration at 15 μM. For all the drug loading tests, *D-*Cys-GQDs/Dox (15 μM/ 200 μM) complexes were incubated with 3T3 exosomes (1.0×10^9^ particles/mL) at room temperature under a static condition. The incubated solution was then washed with 1×PBS buffer under the hold of a 100 kDa cellulose centrifugal filter for multiple times to remove excessive *D-*Cys-GQDs/Dox complex. The change of the fluorescence intensity of the incubated solution was monitored at each washing steps. The intensity of fluorescence decreased as the washing steps increased (**Figure S9a-d**). The fluorescent intensity of *D-*Cys-GQDs/Dox loaded exosomes decreased moderately and kept the retention rate of 30.5% (**Figure S9e**), indicating the strong interaction between *D-*Cys-GQDs/Dox and exosomes. In contrast, relative steep decrease and low retention rate were observed for exosomes loaded by *L--*Cys*-*GQDs/Dox (24.4%), *R-*Cys*-*GQDs/Dox (21.9%), free-Dox (11.5%) and sonication method^56^ (17.5%) (**Figure S8e**).

Drug loading via chiral GQDs into exosomes was further examined by confocal fluorescence microscopy by visualizing the co-localization of chiral GQDs (Blue), exosomes (membrane dye in green) and the Dox (Red) (**Figure S10a**). Due to FRET effect, all blue signals corresponding for GQDs variants were quenched by the attachment of Dox, thus showed relatively low intensity of signal in the confocal images (**Figure S10a**). Thus, the permeation of chiral-GQDs/Dox complex results were analyzed using the Dox channel (Red) (**Figure 5c** and **Figure S10b**). Based on the definition of the encapsulation efficiency in liposome formulation,^57^ drug loading efficiency of exosome is defined to be the percentage of active exosomes that successfully encapsulate drugs(**See Method**). Similar to the permeation efficiency of *D-*Cys-GQDs into exosomes, loading efficiency of *D*-Cys-GQDs/Dox into exosomes (66.7 ± 9.5%) was significantly higher than that of *L*-Cys-GQDs/Dox (18.3 ± 6.7%). Meanwhile, loading efficiency of *R*-Cys-GQDs/Dox (15.2 ± 6.3%) is the lowest among all samples (**Figure 5d**). In addition, exosomes loaded with Dox via traditional sonication approach was prepared as a control. The low loading efficiency of traditional sonication (14.5 ± 7.9%) was reflected with relative low density of Dox signal (Red) on confocal fluorescence microscopy (**Figure 5c**). Most importantly, exosomes retained integrity without significant change in size after drug loading by *D-*Cys-GQDs. In contrast, the physical properties of exosomes were altered by sonication significantly, as shown in TEM and NTA (**Figure S11a, b**). These results demonstrated that drug loading into exosomes facilitated by chiral GQDs had significantly high efficiency and maintained the integrity of exosomes.

To evaluate whether *D*-Cys-GQDs/Dox loaded exosomes (Exo-Dox) can be uptake by the cells, we treated 3T3 cells with Exo-Dox *in vitro* for 12 h and compared it to the control group treated with free Dox. The presence of Dox signals (Red) in the confocal images demonstrated that Dox molecules were delivered successfully into the cells and mainly accumulated in the nucleus (**Figure 5e**). Meanwhile, *D-*Cys-GQDs (Blue) were distributed mostly in the cytosol of cells, indicating the release of Dox from *D-*Cys-GQDs/Dox complex.

We then analyzed the ability of Exo-Dox to inhibit cancer cell proliferation *in vitro*. Human hepatocellular carcinoma cells (HepG2) and cervical carcinoma cells (Hela) were treated with Exo-Dox for 24 h, while cells without treatment, treated with control exosomes, and *D*-Cys-GQDs mixed with free Dox are negative controls. Cell viability of all samples was measured by CCK-8 assay. Exo-Dox inhibited cell proliferation by 44.2 ± 9.2% for HepG2 cells and by 27.4 ± 6.5% for Hela cells, comparable to free Dox (HepG2: 28.3 ± 5.2% and Hela: 27.7 ± 5.0%) (**Figure 5f**). No significant inhibition of cell growth was observed in control samples treated with exosome and *D-*Cys-GQDs, indicating low or no toxicity associated with exosomes or *D-*Cys-GQDs. Dox-loaded exosomes by sonication method were not included in the cell viability tests due to low yield of loaded exosomes that was caused by the damage of physical structure of exosomes by sonication (**Figure S11a, b**).

### Chiral GQD based siRNA loading in exosome for gene therapy

siRNA therapeutics are promising treatment for viral infections, hereditary disorders and cancers.^58^ However, it is still challenging in translational applications due to their the poor intracellular uptake and limited stability of siRNA in blood stream. When siRNA is administered intravenously, it is readily digested by nucleases and largely cleared from the kidney glomeruli before reaching the diseased organs. Exosomes have been invested as nanocarriers for siRNA encapsulation to overcome this challenge due to the high stability in circulation^59^ and efficient cellular uptake^60^ compared with liposome. However, the efficacy of siRNA loading in the exosome is relatively low^7^ due to these nucleotides being relatively large and cannot diffuse into the exosome spontaneously.^61^ Moreover, current methods, such as the usage of transfection reagents and viral transduction-base strategies, may affect the function of exosomes and the pathogenicity and teratogenicity of the viruses may be preserved and inherited in exosomes, resulting in safety risks. ^62,63^

Similar to enhanced Dox loading into exosome through *D-*Cys-GQDs, we utilized *D-*Cys-GQDs to load the siRNA into 3T3 exosomes. The aromatic surface of GQDs can offer an excellent capability to immobilize genomic substant drugs by π-π stacking.^64^ For demonstration of siRNA loading, a nucleic acid sequence (GUGCAAUGAGGGACCAGUA) labeled with red dye ROX (carboxy-X-rhodamine) was tested first. The density of the siRNA on the *D-*Cys*-*GQDs surface was determined as 2 siRNAs per GQD by the same FRET assay (**Figure S12**) as the method for Dox. The optimized condition for loading was chiral-Cys-GQDs/siRNA complex (15 μM / 30 μM) determined by the saturation point of siRNA attachment on *D*-Cys-GQDs for the following loading investigation. The loading of *D-*Cys-GQDs/siRNA in exosomes was confirmed with confocal fluorescence microscopy by the colocalization signals of the labeled siRNA (Red) and exosome (Green, Cellmask green plasma membrane stain) (**Figure 6a**). Furthermore, we visualized *D-*Cys*-*GQDs/siRNA complex loaded exosome through TEM to confirm the successful loading. TEM images showed colocalization of *D-*Cys-GQDs/siRNA and exosomes with black dots in sight of individual exosome compared with the bare one (**Figure 6b**). The loading efficiencies were analyzed using the same method as previous sessions based on the red signal of siRNA (**Figure S13a**). Compared with free siRNA that rarely entered exosomes, the loading efficiencies of *D-*Cys-GQDs/siRNA, *R*-Cys-GQDs/siRNA and *L-*Cys-GQDs/siRNA were 64.1 ± 16.5%%, 17.8 ± 7.4%, and 7.1 ± 4.2% **(Figure 6c** and **Figure S13a**). In addition, the size of siRNA-loaded exosomes remained similar to unloaded exosomes shown in NTA (**Figure S13b**). The results demonstrated that siRNAs were successfully loaded in the exosome by chiral GQDs, and the exosomes remained integrity after the loading procedure.

**Figure 6.**
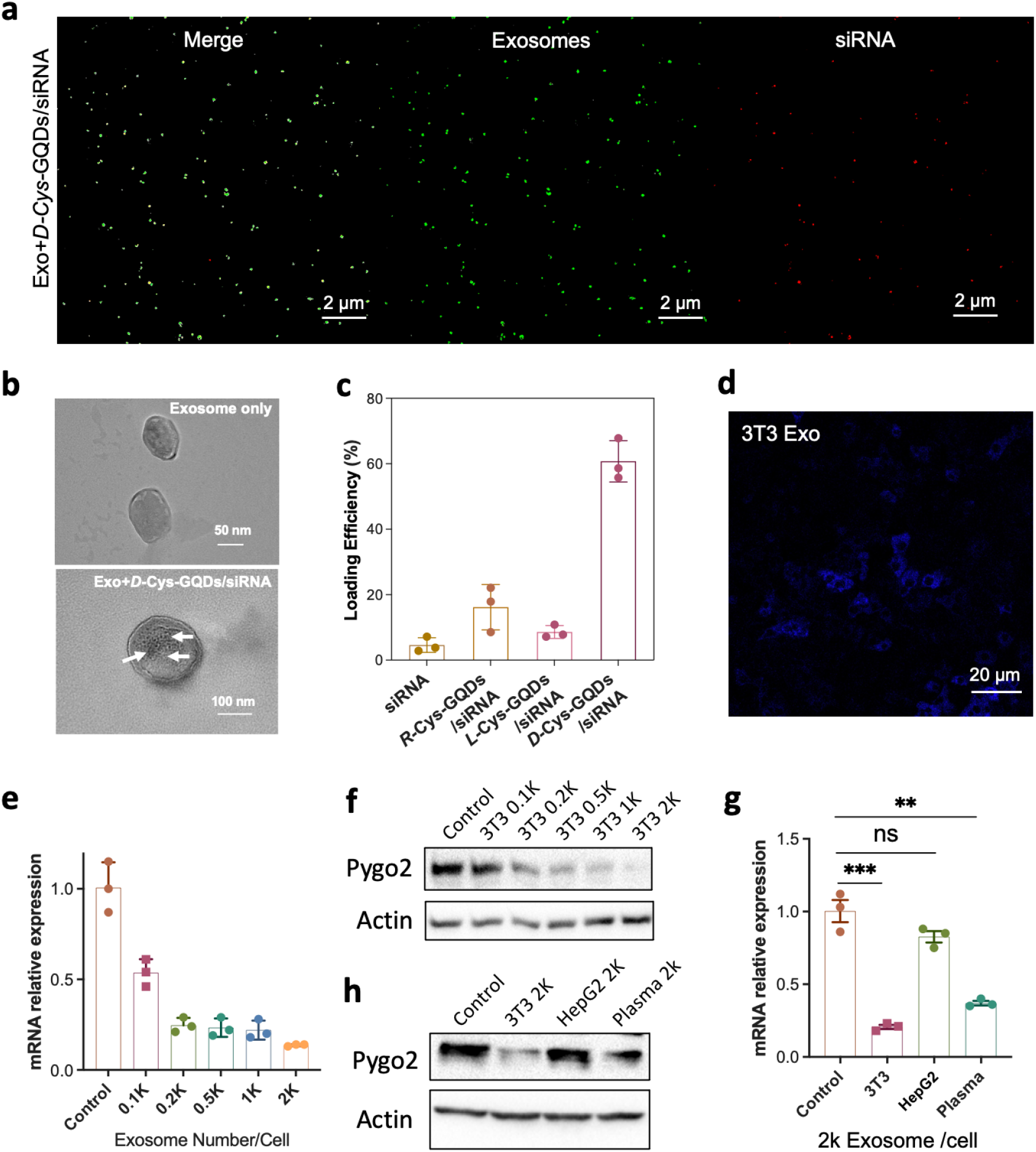
siRNA exosomes loaded by *D*-Cys-GQDs for gene therapy. (a) Confocal images of siRNA (red) loaded exosomes (green) with facilitation of *D-*Cys-GQDs, (b) TEM demonstration of permeation of siRNA with the assistance of *D-*Cys-GQDs, (c) Loading efficiency of chiral Cys-GQDs/siRNA complex into exosomes, (d) Confocal imaging of DU145 cell lines treated with 3T3 exosome for 48h (Scale bars: 20 μm), (e-f) Pygo2 mRNA level(e) and protein level (f) in DU145 cells with the treatment of 3T3 exosome loaded with *D-*Cys-GQDs/Pygo2 siRNA at different density for 48h. (g-h) Comparison of Pygo2 silencing efficiency after treated with *D-*Cys-GQDs/Pygo2 siRNA loaded 3T3, HepG2 or plasma exosome for 48h through qPCR(g) and western blot(h).

To determine whether siRNA loaded exosome by *D-*Cys-GQDs are effective and efficient for cancer treatment, we chose newly identified oncogene Pygo2 in prostate cancer as target. Pygo2 is a chromatin effector and has been reported to have overexpression in prostate, ovarian, breast, cervical, hepatic, lung, intestinal, and brain cancers.^65^ We recently also found that Pygo2 played an essential role in immunosuppressive tumor microenvironment regulation and inducing the resistance of prostate cancer to immunotherapy.^66^ Therefore, targeting Pygo2 has very good implication in clinical treatment for many types of cancer. We have demonstrated the knock-down of gene using commercially available Pygo2 (Pygopus homolog 2) siRNA.^67^ Pygo2 has good efficiency on silencing and has been used for gene delivery application.^67^ Meanwhile, DU145 is a widely used human prostate cancer cell line with a high expression of Pygo2. Thus, we assessed *D-*Cys-GQDs/siRNA loaded exosomes using DU145 cell culture *in vitro*. First, we tested the uptake of the isolated 3T3 exosomes by DU145 cell line. A strong fluorescent signal (Blue: GQDs) was observed in the cytoplasm of DU145 cells incubated with *D-*Cys-GQDs loaded (3T3) exosomes for 48 hours (**Figure 6d**). Efficiencies of gene knock-down by *D-*Cys-GQDs/siRNA loaded exosomes were confirmed with qRT-PCR and Western blotting. For 3T3 exosomes (100 to 2000 Exo/cell), it showed significantly reduction of Pygo2 mRNA levels by 45%∼ 80% with a mean value of 62% (**Figure 6e**). The expression of Pygo2 protein levels in DU145 cells were also silenced by *D-*Cys-GQDs/siRNA loaded (3T3) exosomes by 22%∼ 91%, compared to the control group (**Figure 6f**). In addition, *D-*Cys-GQDs/siRNA induced a dose-dependent decrease for both mRNA and protein expressions. To reflect loading reliability of *D-*Cys-GQDs, exosomes isolated from two other sources, HepG2 cell culture medium and healthy human plasma, were tested as controls. Similarly, *D-*Cys-GQDs/siRNA loaded plasma exosomes reduced expression of mRNA by 63.3 ± 7.8% (**Figure 6g**) and expression of protein by 50% (**Figure 6h)**. HepG2 exosome showed no obvious reduction of mRNA and Pygo2 protein, which contributed to the fact that the exosome uptake capabilities were different depending on the recipient cell types.^68^ An overall 60%–80% knock-down of the target gene and higher than 60% inhibition of Pygo2 protein, indicating the efficacy of *D-*Cys-GQDs/siRNA loaded exosomes to the target cells. Overall, *D-*Cys-GQDs can facilitate loading genes such as siRNA into exosomes without membrane damage of exosomes and the loaded exosomes can successfully mediate silencing of target genes with high efficiency.

## Conclusion

In summary, we have developed an exogenous drug-agnostic chiral GQDs exosome-loading platform, based on chirality matching with the exosome lipid bilayer. Both hydrophobic and hydrophilic chemical and biological drugs can be functionalized or adsorbed onto GQDs by π–π stacking and van der Waals interactions. By tuning the ligands and GQD size to optimize its chirality, we demonstrate significantly high drug loading efficiency for Dox and siRNA into exosomes by *D-*Cys-GQDs at an optimal concentration, compared to other reported exosome loading techniques.^14^ The drug loaded exosomes by *D*-Cys-GQDs are shown to be effective in killing cancer cells, the knock-down of the target gene, and the inhibition of mRNA and relative protein expression levels. Thus, chiral GQD enhanced drug loading is a promising generic and scalable drug loading technique that can enable high-throughput production of therapeutic exosomes for clinical applications.

### Method and Experimental Section Isolation and characterization of exosomes

The Hep-G2 and 3T3 cells were purchased from the American Type Culture Collection (ATCC) and propagated in minimum essential medium (MEM, Corning Incorporated, Corning, NY, USA) supplemented with 10% FBS (exosome-depleted) and antibiotics. All cells were maintained in 5% CO_2_ at 37 °C and the cell culture medium (CM) was collected for 24 hours. Then, the CM was centrifuged at 500g for 10 min, 2000g for 10 min, and 12,000g for 30 min to remove cells and cell debris. The supernatants were pelleted by ultracentrifugation at 100,000g for 80 min. However, we found the yield of ultracentrifugation to be too low to produce sufficient exosomes for loading, to the extent that pellets were not found for some cell media samples. Instead, we adopted an ultrafiltration method based on our earlier work. The nanopores of ion-track membranes are etched into conic pores^69^ to allow high-throughput isolation, enrichment and purification with minimum loss of the exosomes.^70, 71^ The data reported are mostly from this Asymmetric Nanopore Membrane (ANM) ultrafiltration technique with a flow rate of 20 ml/hour. A typical NTA characterization of ANM filtered exosome is seen in Fig S1b in Supp Material. Exosome were dissolved with PBS buffer and stored at −80°C until use.

Exosomal markers CD63[1:1000; Cell Signaling Technology (CST), USA], and Alix (1:1000; Abcam, UK) and the ER marker calnexin (1:1000; Abcam, UK) were detected by Western blot. The size distribution and zeta potential of N-Ex were measured by the NanoSight LM10 system (NTA, UK). The PDI of N-Ex was tested by DLS (Malvern Instruments, UK).

Nanoparticle tracking analysis (NTA) measurements were performed with a NanoSight NS300 (NanoSight Ltd., UK) using purified exosomes (100uL in 1 mL PBS buffer). The mean exosome size distribution (modal hydrodynamic diameter in nm) and exosome concentration (number of EVs enriched from 1 mL of sample in particles/mL) were captured and analyzed with the NTA 3.3 Analytical Software Suite. All procedures were performed at room temperature.

The morphology of exosome was identified by TEM (JEOL 2011) and AFM (Park XE7, Korea). Its structure was further characterized by electron microscopy (EM). Purified exosomes were resuspended in PBS and fixed with 2% paraformaldehyde for 30 min at room temperature. Eight microliters of mixture were then dropped onto EM grids that had been pretreated with UV light to reduce static electricity. After drying for 30 min, exosomes were stained twice (6 min each) with 1% uranyl acetate. The dried grids were examined using an HT7700 (JEOL 2011) transmission electron microscope (TEM) at 120 kV.

### GQD and Chiral GQD Synthesis

The carbon nanofibers (100 nm), *L/D*-Cysteine, sulfuric acid, nitric acid, N-(3-Dimethylaminopropyl)-N’-ethylcarbodiimide hydrochloride (EDC, 191.7 g/mol) and N-Hydroxysulfosuccinimide sodium salt (Sulfo-NHS, 217.13 g/mol) were purchased from Sigma-Aldrich. The GQDs were synthesized by a modified protocol from our previous report.^19, 72^ Briefly, 0.4 g of carbon nanofibers was dispersed into a 40 mL mixture of sulfuric acid and nitric acid (3:1, v/v) and sonicated for 2-h. The mixture was mechanically stirred for 6-h at room temperature and followed by being heated to 120 °C to continuously react for 10-h. After the reaction, the mixture solution was cooled and diluted with ice DI water and adjusted pH to 8 by adding sodium hydroxide. Then, the GQDs was purified with 3 days dialysis and the final concentration of GQDs was 1 mg/mL. In order to impart chirality to the GQDs, the carboxylic group of GQDs was connected with the amine group of *L-*(or *D-*)cysteine by the EDC/NHS method.^55, 73^ Briefly, a solution of EDC (20 μL, 100 mM) was added into 2 mL of GQD (100 μM) solution. After 10 min stirring, the 40 μL of Sulfo-NHS (100 mM) was added to the solution, and it was sonicated for 40 min under an ice−water bath.^74^ The resulting mixture was treated by a 1kDa centrifuge tube and rinsed for three times to remove excess EDC and sulfo-NHS. Finally, 40 μL of *L-* (or *D-*) cysteine (100 mM) was added into the GQD-NHS ester, and the mixture was stirred for 2-h. The surplus L/D-form of cysteine was removed by a dialysis membrane (1 kDa, Fisher Scientific). Relatively large size GQDs were obtained with average size tunable by reaction time.^75^ After 2 h of reaction under the same condition (sulfuric acid and nitric acid were 3:1 in v/v), the size of GQDs were found to be around 3-90 nm (mean value: 50 nm). The solution was separated and cut off by using two sizes of nanoporous membrane (18 and 50 nm). The small size GQDs (< 18 nm) in this bench was discard. The middle size (18 ∼ 50 nm) and large size (50 ∼ 90 nm) in this bench was collected and then they were modified with *L/D*-cysteine using the same EDC/NHS method. The certain size was characterized in the Figure 2 by TEM.

The chiroptical activity of the dispersions was measured by CD spectroscopy (J-1700, JASCO), and the chemical reaction progress was monitored by FT-IR spectroscopy (FT/IR-6300, Jasco) and Raman spectroscopy (NRS-5100, Jasco). The absorbance of chiral GQD was analyzed by UV/vis spectroscopy (Agilent, 89090A). The fluorescence property of chiral GQDs was characterized by Infinite M1000 plate reader (Tecan Group). The morphology of chiral GQDs was observed by Transmission Electron Microscopy (TEM) (JEOL 2011). Their surface potential was analyzed by a Zetasizer (Malvern Instruments, Nano ZS).

### Cell Cultures and Viability Assays

3T3 and Hepatocellular carcinoma human cells (HepG2) (ATCC, VA) were maintained with Eagle’s minimum essential medium (EMEM) supplemented with 10% fetal bovine serum (FBS) and 1% penicillin−streptomycin (ATCC) in a humidified incubator (MCO-15AC, Sanyo) at 37 °C in which the CO2 level was maintained at 5% before seeding. All of the medium was filtered using 0.22 μ SteriCup filter assembly (Millipore, USA) and stored at 4 °C for no longer than 2 weeks. For cells incubated with the siRNA-exosomes, and control exosomes, the cells were cultured overnight to allow attachment in a 96-well plate and confocal dishes, washed with FBS-free EMEM, and then incubated with exosome analytes at 37 °C for 1 h in FBS-free medium. After incubation with difference windows, the cells were washed repeatedly with sterilized PBS and maintained in culture medium before further analysis.

Cell viability was assessed using a Cell Counting Kit-8 (CCK-8 assay, Enzo). In brief, 3T3 and HepG2 cells were seeded into a 96-well flat culture plate (Corning). After being cultured overnight, the cells were washed with FBS-free EMEM and incubated with a specific concentration of exosome and exosomal relative in FBS-free medium at 37 °C for 12, 24, 36, 48-h. The cells were then washed three times with sterilized PBS and incubated with fresh medium containing 10% FBS overnight. The cells were then washed with PBS, and FBS-free EMEM (500 mL) before adding assay reagent. After incubation for 20 min at 37 °C, the cell proliferation and cytotoxicity were measured by absorbance at 460 nm. The background was measured by 3T3 and HepG2 cells cultured in FBS-free EMEM only.

### Loading therapeutic cargo

To load the exosomes with Dox, 150 μL of purified exosomes and 50 μL of complex (40 μM *D-*Cys-GQDs: 560 μM Dox) were gently mixed at 4 °C and the mixture was incubated at room temperature for 20 min to ensure the fully permission. To load the exosomes with siRNA, 150 μL of purified exosomes and 50 μL of complex (40 μM *D*-Cys-GQDs: 80 μM siRNA) were gently mixed at 4 °C and the mixture was incubated at 37 °C for 20 min to ensure the fully permeation. Recovery was assessed by analysis of confocal image as described above. Exosomes were then washed with cold PBS four times under the supporting of 100kDa centrifuge tube to remove unincorporated free *D-*Cys-GQDs/siRNA. The exosomes loaded with siRNA were quantified for the encapsulated genomic drug by detecting the intrinsic fluorescence of labeled siRNA using Infinite M1000 plate reader (Tecan Group) at 608 nm with excitation at 588 nm and image analysis of confocal microscope under the excitation at 561 nm.

### Determination of the permeation efficiency of chiral GQDs

The permeation efficiency of GQDs in exosomes were indirectly determined by statistically analyzing the count of blue fluorescent light-up exosomes (caused by permeation) under confocal microscopy over the total concentration of exosome after loading. In brief, 5μL loaded exosome sample was fully covered with 18 mm × 18 mm cover glasses (Corning®, square, No. 1) and scanned under 100× Objective with normal field size 150 μm × 150 μm. Four Z-stacks of images were captured through random domains. Each Z-stack contained around 20∼30 images from presenting to disappearing fluorescent dots with a step size of 0.125 μm. Then, captured images were analyzed using ImageJ. The total fluorescent exosome particles (TFEPs) were counted by settings with manually adjusted thresholds and matching the size of exosomes. Colocalized fluorescent exosome particles (CFEPs) of Z-stack images was counted based on centers of mass-particles coincidence by using the JACOPx Plugin. The Permeation efficiency into exosomes was calculated using the following formula:

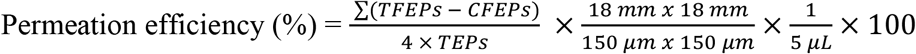

where TECs is the total exosome concentration (particles/mL) that is measured by NTA.

### Determination of loading efficiency of drug in exosomes

While the success of the drug loading procedure was mostly reported in the form of loading efficiency (the percentage of total available drug that has been encapsulated within EVs) or loading capacity (the amount of drug loaded per mass of particles),^76^ it is only applicable for reflecting the concentration of active drugs and cannot realistically be applied for defining reproducible protocols of the exogenous loading of exosomes.^14^ By taking the encapsulation efficiency of liposome formulation,^57^ loading efficiency of exosome, which measures the percentage of active exosomes that successfully encapsulate drugs, was used in the design of exosome-based drug delivery systems. Same as the GQDs permeation, the loading efficiency of Dox and siRNA in exosomes were indirectly determined by statistically analyzing the count of red fluorescent light-up exosomes (caused by loading) under confocal microscopy over the total concentration of exosome after loading. The loading efficiency into exosomes was calculated using the following formula:

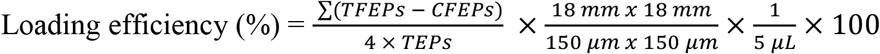

where TECs is the total exosome concentration (particles/mL) that is measured by NTA.

## Supporting information

Supporting Information_Chiral Graphene Quantum Dots (GQDs) Enhanced Drug Delivery in Exosomes

## ASSOCIATE CONTENT

### Supporting Information

Additional figures, including characterization of 3T3 exosome, GQDs and chiral GQDs; penetration study of chiral GQDs on exosome; TEM images of exosomes treated with *D-*Cys-GQDs; confocal images of PHK26 labelled exosomes; UV–vis absorption and zeta-potential (ζ) of the *L-/D-*Arg-GQDs and *L-/D-*Trp-GQDs; effect of *D-*Cys-GQDs concentration on fluorescence intensity and quenching efficiency of Dox, and the profile of siRNA loaded exosomes.

## Author Contributions

Yichun Wang and Youwen Zhang conceptualized the study. Youwen Zhang designed and carried out the experiments. Yini Zhu did the in vitro test, quantified the protein expression and mRNA levels. Gaeun Kim did biological experiments, compared the exosome loading with sonication method and isolated exosome by ultracentrifuge. Ceming Wang isolated exosome by ANM. Runyao Zhu took the TEM images and helped to synthesize chiral GQDs, Hsueh-Chia Chang and Yichun Wang supervised all the work, administrated the project, and acquired the funding. Youwen Zhang prepared the manuscript, and all authors contributed to data interpretation, discussions, and writing.

## Notes

The authors declare no competing financial interests.

## Acknowledgments

This work was financially supported by American Cancer Society Institutional Research Grant (ACS IRG-17-182-04) at the Harper Cancer Research Institute, the National Science Foundation Industry-University Cooperative Research Center (The Center for Bioanalytic Metrology) and the NIH Common Fund, through the Office of Strategic Coordination/Office of the NIH Director, 1UG3CA241684-01.

## TOC

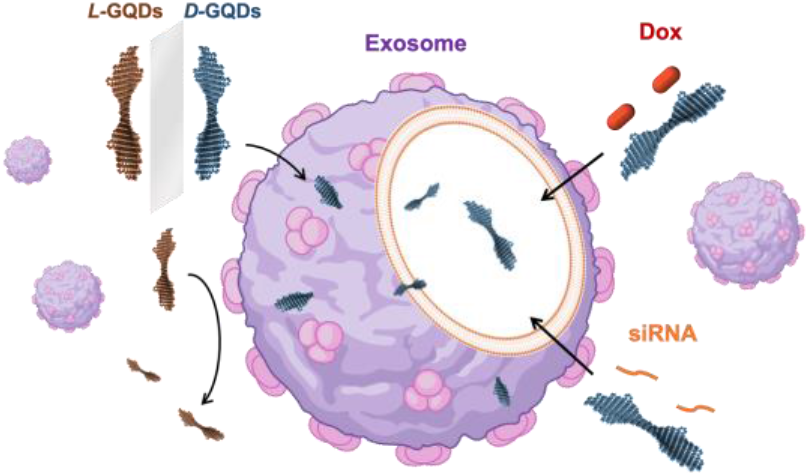

## Reference

1. Tenchov R, Sasso JM, Wang X, Liaw W-S, Chen C-A, Zhou QA. Exosomes-Nature’s Lipid Nanoparticles, a Rising Star in Drug Delivery and Diagnostics. ACS Nano 2022, 16(11): 17802–17846.

2. Fu S, Wang Y, Xia X, Zheng JC. Exosome engineering: Current progress in cargo loading and targeted delivery. NanoImpact 2020, 20: 100261.

3. Kimiz-Gebologlu I, Oncel SS. Exosomes: Large-scale production, isolation, drug loading efficiency, and biodistribution and uptake. J Controlled Release 2022, 347: 533–543.

4. Chen H, Wang L, Zeng X, Schwarz H, Nanda HS, Peng X, et al. Exosomes, a New Star for Targeted Delivery. Front Cell Dev Biol 2021, 9: 751079.

5. Shahabipour F, Barati N, Johnston TP, Derosa G, Maffioli P, Sahebkar A. Exosomes: Nanoparticulate tools for RNA interference and drug delivery. J Cell Physiol 2017, 232(7): 1660–1668.

6. Liu C, Su C. Design strategies and application progress of therapeutic exosomes. Theranostics 2019, 9(4): 1015–1028.

7. Xu M, Yang Q, Sun X, Wang Y. Recent Advancements in the Loading and Modification of Therapeutic Exosomes. Frontiers in Bioengineering and Biotechnology 2020, 8.

8. Wang J, Chen D, Ho EA. Challenges in the development and establishment of exosome-based drug delivery systems. J Controlled Release 2021, 329: 894–906.

9. Morshedi Rad D, Alsadat Rad M, Razavi Bazaz S, Kashaninejad N, Jin D, Ebrahimi Warkiani M. A Comprehensive Review on Intracellular Delivery. Adv Mater (Weinheim, Ger) 2021, 33(13): 2005363.

10. Sensale S, Peng Z, Chang H-C. Acceleration of DNA melting kinetics using alternating electric fields. The Journal of Chemical Physics 2018, 149(8): 085102.

11. Slouka Z, Senapati S, Chang H-C. Microfluidic Systems with Ion-Selective Membranes. Annu Rev Anal Chem 2014, 7(1): 317–335.

12. Fuhrmann G, Serio A, Mazo M, Nair R, Stevens MM. Active loading into extracellular vesicles significantly improves the cellular uptake and photodynamic effect of porphyrins. J Controlled Release 2015, 205: 35–44.

13. Herrmann IK, Wood MJA, Fuhrmann G. Extracellular vesicles as a next-generation drug delivery platform. Nat Nanotechnol 2021, 16(7): 748–759.

14. Rankin-Turner S, Vader P, O’Driscoll L, Giebel B, Heaney LM, Davies OG. A call for the standardised reporting of factors affecting the exogenous loading of extracellular vesicles with therapeutic cargos. Adv Drug Deliv Rev 2021, 173: 479–491.

15. O’Loughlin AJ, Mäger I, de Jong OG, Varela MA, Schiffelers RM, El Andaloussi S, et al. Functional Delivery of Lipid-Conjugated siRNA by Extracellular Vesicles. Mol Ther 2017, 25(7): 1580–1587.

16. Pomatto MAC, Bussolati B, D’Antico S, Ghiotto S, Tetta C, Brizzi MF, et al. Improved Loading of Plasma-Derived Extracellular Vesicles to Encapsulate Antitumor miRNAs. Mol Ther Methods Clin Dev 2019, 13: 133–144.

17. Haney MJ, Klyachko NL, Zhao Y, Gupta R, Plotnikova EG, He Z, et al. Exosomes as drug delivery vehicles for Parkinson’s disease therapy. J Control Release 2015, 207: 18–30.

18. Chen P, Yue H, Zhai X, Huang Z, Ma G-H, Wei W, et al. Transport of a graphene nanosheet sandwiched inside cell membranes. Science Advances 2019, 5(6): eaaw3192.

19. Suzuki N, Wang Y, Elvati P, Qu Z-B, Kim K, Jiang S, et al. Chiral Graphene Quantum Dots. ACS Nano 2016, 10(2): 1744–1755.

20. Liu C, Elvati P, Majumder S, Wang Y, Liu AP, Violi A. Predicting the Time of Entry of Nanoparticles in Lipid Membranes. ACS Nano 2019, 13(9): 10221–10232.

21. Yeom J, Guimaraes PPG, Ahn HM, Jung B-K, Hu Q, McHugh K, et al. Chiral Supraparticles for Controllable Nanomedicine. Adv Mater (Weinheim, Ger) 2020, 32(1): 1903878.

22. Baimanov D, Wang J, Zhang J, Liu K, Cong Y, Shi X, et al. In situ analysis of nanoparticle soft corona and dynamic evolution. Nature Communications 2022, 13(1): 5389.

23. Mitchell MJ, Billingsley MM, Haley RM, Wechsler ME, Peppas NA, Langer R. Engineering precision nanoparticles for drug delivery. Nat Rev Drug Discovery 2021, 20(2): 101–124.

24. Yeom J, Guimaraes PPG, Ahn HM, Jung BK, Hu Q, McHugh K, et al. Chiral Supraparticles for Controllable Nanomedicine. Adv Mater 2020, 32(1): e1903878.

25. Chen F, Gao W, Qiu X, Zhang H, Liu L, Liao P, et al. Graphene quantum dots in biomedical applications: Recent advances and future challenges. Frontiers in Laboratory Medicine 2017, 1(4): 192–199.

26. Hai X, Feng J, Chen X, Wang J. Tuning the optical properties of graphene quantum dots for biosensing and bioimaging. Journal of Materials Chemistry B 2018, 6(20): 3219–3234.

27. Tian P, Tang L, Teng KS, Lau SP. Graphene quantum dots from chemistry to applications. Materials Today Chemistry 2018, 10: 221–258.

28. Baimanov D, Wu J, Chu R, Cai R, Wang B, Cao M, et al. Immunological Responses Induced by Blood Protein Coronas on Two-Dimensional MoS2 Nanosheets. ACS Nano 2020, 14(5): 5529–5542.

29. Razmi H, Mohammad-Rezaei R. Graphene quantum dots as a new substrate for immobilization and direct electrochemistry of glucose oxidase: Application to sensitive glucose determination. Biosensors and Bioelectronics 2013, 41: 498–504.

30. Li Z, Fan J, Tong C, Zhou H, Wang W, Li B, et al. A smart drug-delivery nanosystem based on carboxylated graphene quantum dots for tumor-targeted chemotherapy. Nanomedicine (Lond) 2019, 14(15): 2011–2025.

31. Ahmadi-Kashani M, Dehghani H, Zarrabi A. A biocompatible nanoplatform formed by MgAl-layered double hydroxide modified Mn3O4/N-graphene quantum dot conjugated-polyaniline for pH-triggered release of doxorubicin. Materials Science and Engineering: C 2020, 114: 111055.

32. Skotland T, Sandvig K, Llorente A. Lipids in exosomes: Current knowledge and the way forward. Prog Lipid Res 2017, 66: 30–41.

33. Abels ER, Breakefield XO. Introduction to Extracellular Vesicles: Biogenesis, RNA Cargo Selection, Content, Release, and Uptake. Cellular and Molecular Neurobiology 2016, 36(3): 301–312.

34. Li G, Luican A, Lopes dos Santos JMB, Castro Neto AH, Reina A, Kong J, et al. Observation of Van Hove singularities in twisted graphene layers. Nat Phys 2010, 6(2): 109–113.

35. Zhang S, Song A, Chen L, Jiang C, Chen C, Gao L, et al. Abnormal conductivity in low-angle twisted bilayer graphene. Science Advances 2020, 6(47): eabc5555.

36. Noor-Ul-Ain, Eriksson MO, Schmidt S, Asghar M, Lin P-C, Holtz PO, et al. Tuning the Emission Energy of Chemically Doped Graphene Quantum Dots. Nanomaterials 2016, 6(11): 198.

37. Jin SH, Kim DH, Jun GH, Hong SH, Jeon S. Tuning the Photoluminescence of Graphene Quantum Dots through the Charge Transfer Effect of Functional Groups. ACS Nano 2013, 7(2): 1239–1245.

38. Huang B, Bates M, Zhuang X. Super-resolution fluorescence microscopy. Annu Rev Biochem 2009, 78: 993–1016.

39. Aiello CD, Abendroth JM, Abbas M, Afanasev A, Agarwal S, Banerjee AS, et al. A Chirality-Based Quantum Leap. ACS Nano 2022, 16(4): 4989–5035.

40. Russ KA, Elvati P, Parsonage TL, Dews A, Jarvis JA, Ray M, et al. C60 fullerene localization and membrane interactions in RAW 264.7 immortalized mouse macrophages. Nanoscale 2016, 8(7): 4134–4144.

41. Guo R, Mao J, Yan L-T. Computer simulation of cell entry of graphene nanosheet. Biomaterials 2013, 34(17): 4296–4301.

42. Dallavalle M, Calvaresi M, Bottoni A, Melle-Franco M, Zerbetto F. Graphene Can Wreak Havoc with Cell Membranes. ACS Appl Mater Interfaces 2015, 7(7): 4406–4414.

43. Li Y, Wang X, Miao J, Li J, Zhu X, Chen R, et al. Chiral Transition Metal Oxides: Synthesis, Chiral Origins, and Perspectives. Adv Mater (Weinheim, Ger) 2020, 32(41): 1905585.

44. Vázquez-Nakagawa M, Rodríguez-Pérez L, Martín N, Herranz MÁ. Supramolecular Assembly of Edge Functionalized Top-Down Chiral Graphene Quantum Dots. Angewandte Chemie International Edition 2022, 61(43): e202211365.

45. Kuznetsova VA, Mates-Torres E, Prochukhan N, Marcastel M, Purcell-Milton F, O’Brien J, et al. Effect of Chiral Ligand Concentration and Binding Mode on Chiroptical Activity of CdSe/CdS Quantum Dots. ACS Nano 2019, 13(11): 13560–13572.

46. Du Y-X, Zhou L-J, Guo J-G. The influence of strain range, size and chiral on mechanical properties of graphene: Molecular dynamics insights. Nanomaterials and Nanotechnology 2022, 12: 18479804221110023.

47. Monti D, Venanzi M, Stefanelli M, Sorrenti A, Mancini G, Di Natale C, et al. Chiral Amplification of Chiral Porphyrin Derivatives by Templated Heteroaggregation. J Am Chem Soc 2007, 129(21): 6688–6689.

48. Ghanbari N, Salehi Z, Khodadadi AA, Shokrgozar MA, Saboury AA, Farzaneh F. Tryptophan-functionalized graphene quantum dots with enhanced curcumin loading capacity and pH-sensitive release. Journal of Drug Delivery Science and Technology 2021, 61: 102137.

49. Eda G, Lin Y-Y, Mattevi C, Yamaguchi H, Chen H-A, Chen I-S, et al. Blue Photoluminescence from Chemically Derived Graphene Oxide. Adv Mater (Weinheim, Ger) 2010, 22(4): 505–509.

50. Nguyen M-K, Kuzyk A. Reconfigurable Chiral Plasmonics beyond Single Chiral Centers. ACS Nano 2019, 13(12): 13615–13619.

51. Yao H, Miki K, Nishida N, Sasaki A, Kimura K. Large Optical Activity of Gold Nanocluster Enantiomers Induced by a Pair of Optically Active Penicillamines. J Am Chem Soc 2005, 127(44): 15536–15543.

52. Ogawa Y. Electron microdiffraction reveals the nanoscale twist geometry of cellulose nanocrystals. Nanoscale 2019, 11(45): 21767–21774.

53. Wallbrecher R, Ackels T, Olea RA, Klein MJ, Caillon L, Schiller J, et al. Membrane permeation of arginine-rich cell-penetrating peptides independent of transmembrane potential as a function of lipid composition and membrane fluidity. J Controlled Release 2017, 256: 68–78.

54. Jang SC, Kim OY, Yoon CM, Choi D-S, Roh T-Y, Park J, et al. Bioinspired Exosome-Mimetic Nanovesicles for Targeted Delivery of Chemotherapeutics to Malignant Tumors. ACS Nano 2013, 7(9): 7698–7710.

55. Zhang Y, Chen X, Yuan S, Wang L, Guan X. Joint Entropy-Assisted Graphene Oxide-Based Multiplexing Biosensing Platform for Simultaneous Detection of Multiple Proteases. Anal Chem 2020, 92(22): 15042–15049.

56. Haney MJ, Zhao Y, Jin YS, Li SM, Bago JR, Klyachko NL, et al. Macrophage-Derived Extracellular Vesicles as Drug Delivery Systems for Triple Negative Breast Cancer (TNBC) Therapy. Journal of Neuroimmune Pharmacology 2020, 15(3): 487–500.

57. Yamamoto E, Miyazaki S, Aoyama C, Kato M. A simple and rapid measurement method of encapsulation efficiency of doxorubicin loaded liposomes by direct injection of the liposomal suspension to liquid chromatography. Int J Pharm 2018, 536(1): 21–28.

58. Kanasty R, Dorkin JR, Vegas A, Anderson D. Delivery materials for siRNA therapeutics. Nat Mater 2013, 12(11): 967–977.

59. Chan M-H, Chang Z-X, Huang C-YF, Lee LJ, Liu R-S, Hsiao M. Integrated therapy platform of exosomal system: hybrid inorganic/organic nanoparticles with exosomes for cancer treatment. Nanoscale Horizons 2022, 7(4): 352–367.

60. Lu M, Zhao X, Xing H, Xun Z, Zhu S, Lang L, et al. Comparison of exosome-mimicking liposomes with conventional liposomes for intracellular delivery of siRNA. Int J Pharm 2018, 550(1): 100–113.

61. Luan X, Sansanaphongpricha K, Myers I, Chen H, Yuan H, Sun D. Engineering exosomes as refined biological nanoplatforms for drug delivery. Acta Pharmacol Sin 2017, 38(6): 754–763.

62. Zhang Y, Liu Y, Liu H, Tang WH. Exosomes: biogenesis, biologic function and clinical potential. Cell & Bioscience 2019, 9(1): 19.

63. Chen H, Wang L, Zeng X, Schwarz H, Nanda HS, Peng X, et al. Exosomes, a New Star for Targeted Delivery. Frontiers in Cell and Developmental Biology 2021, 9.

64. Ling K, Jiang H, Li Y, Tao X, Qiu C, Li F-R. A self-assembling RNA aptamer-based graphene oxide sensor for the turn-on detection of theophylline in serum. Biosensors and Bioelectronics 2016, 86: 8–13.

65. Li Q, Li Y, Gu B, Fang L, Zhou P, Bao S, et al. Akt Phosphorylates Wnt Coactivator and Chromatin Effector Pygo2 at Serine 48 to Antagonize Its Ubiquitin/Proteasome-mediated Degradation. J Biol Chem 2015, 290(35): 21553–21567.

66. Talla SB, Brembeck FH. The role of Pygo2 for Wnt/ß-catenin signaling activity during intestinal tumor initiation and progression. Oncotarget 2016, 7(49).

67. Lu X, Pan X, Wu CJ, Zhao D, Feng S, Zang Y, et al. An In Vivo Screen Identifies PYGO2 as a Driver for Metastatic Prostate Cancer. Cancer Res 2018, 78(14): 3823–3833.

68. Murphy DE, de Jong OG, Brouwer M, Wood MJ, Lavieu G, Schiffelers RM, et al. Extracellular vesicle-based therapeutics: natural versus engineered targeting and trafficking. Exp Mol Med 2019, 51(3): 1–12.

69. Wang C, Sensale S, Pan Z, Senapati S, Chang H-C. Slowing down DNA translocation through solid-state nanopores by edge-field leakage. Nature Communications 2021, 12(1): 140.

70. Zhang C, Huo X, Zhu Y, Higginbotham JN, Cao Z, Lu X, et al. Electrodeposited magnetic nanoporous membrane for high-yield and high-throughput immunocapture of extracellular vesicles and lipoproteins. Communications Biology 2022, 5(1): 1358.

71. Wang C, Senapati S, Chang H-C. Liquid biopsy technologies based on membrane microfluidics: High-yield purification and selective quantification of biomarkers in nanocarriers. Electrophoresis 2020, 41(21-22): 1878–1892.

72. Zhang Y, Chen X, Roozbahani GM, Guan X. Graphene oxide-based biosensing platform for rapid and sensitive detection of HIV-1 protease. Anal Bioanal Chem 2018, 410(24): 6177–6185.

73. Zhang Y, Chen X, Roozbahani GM, Guan X. Rapid and sensitive detection of the activity of ADAM17 using a graphene oxide-based fluorescence sensor. Analyst 2019, 144(5): 1825–1830.

74. Zhang Y, Chen X, Wang C, Roozbahani GM, Chang H-C, Guan X. Chemically functionalized conical PET nanopore for protein detection at the single-molecule level. Biosensors and Bioelectronics 2020, 165: 112289.

75. Luo J, Cote LJ, Tung VC, Tan ATL, Goins PE, Wu J, et al. Graphene Oxide Nanocolloids. J Am Chem Soc 2010, 132(50): 17667–17669.

76. Kim J, Shamul JG, Shah SR, Shin A, Lee BJ, Quinones-Hinojosa A, et al. Verteporfin-Loaded Poly(ethylene glycol)-Poly(beta-amino ester)-Poly(ethylene glycol) Triblock Micelles for Cancer Therapy. Biomacromolecules 2018, 19(8): 3361–3370.

