## Supporting Information_Chiral Graphene Quantum Dots (GQDs) Enhanced Drug Delivery in Exosomes for "Chiral Graphene Quantum Dots Enhanced Drug Loading into Exosomes"

**Table of Contents :**

**Figure S1.** Characterizations of GQDs and chiral GQDs ........................................ S-3

**Figure S2.** Characterization of 3T3 exosome........................................................ S-4

**Figure S3.** Penetration study of chiral GQDs on exosome ........................................ S-5

**Figure S4.** TEM images of exosomes treated with *D*-Cys-GQDs .............................. S-6

**Figure S5.** Size effect on permeation of chiral GQDs into exosome .......................... S-7

**Figure S6.** (a) UV–vis absorption and (b) zeta-potential (ζ) of the *L-/D-* Arg-GQDs and *L-/D-* Trp-GQDs........................................................................................ S-8

**Figure S7.** TEM images of damaged exosomes that was treated with *D*-Cys-GQDs at the concentration of 30 μM.......................................................................... S-9

**Figure S8.** Effect of *D*-GQDs concentration on (a) fluorescence intensity and (b) quenching efficiency of Dox, (c) UV−vis absorbance spectra of Dox, *D*-Cys-GQDs, and *D*-Cys-GQDs/Dox complex ........................................................................ S-10

**Figure S9.** Penetration study of chiral Cys-GQDs/Dox on exosome.......................... S-11

**Figure S10.** Confocal image of free Dox, *R-, L-, D-* Cys-GQDs/Dox and sonication treated exosomes........................................................................................... S-12

**Figure S11.** Comparison of Dox loading by *D*-Cys-GQDs and sonication ................ S-13

**Figure S12.** (a) Effect of GQDs concentration on fluorescence intensity and (b) quenching efficiency of 10uM siRNA........................................................................... S-14

**Figure S13.** The profile of siRNA loaded exosomes................................................. S-15

### Additional Methods

#### Isolation and characterization of Exosome:

To investigate the permeability of *D*-GQD into exosomes, the exosomes were harvested from cell cultures of mouse fibroblast cell line (3T3) in vitro. Ultrafiltration and density-gradient ultracentrifugation are often necessary to isolate and purify the exosomes the high shear stress of such isolation procedures often induces protein denaturation and aggregation. For example, low density lipoproteins (LDLs) are known to form aggregates that are in the same 30 to 200 nm size range of exosomes and are difficult to separate from the exosomes. To ensure exosome purity during drug loading by chiral GQD, we employed an ion-track ultrafiltration membrane whose pores have been etched into a conic geometry to reduce shear and fouling by the proteins during their transit. The conic tips of these asymmetric nanopore membrane (ANM)[1] can be etched down to 30 nm, with 3% variation from pore to pore, to prevent exosome transit. Cholesterol assays for a magnetic version of ANM have shown undetectable LDL concentration in the isolated exosomes[2,3]. Although exosomes from serum-free cell culture media are used in our loading experiments, we nevertheless use ANM to purify and enrich our exosomes. The isolated 3T3 exosome displayed a cup-shaped morphology (**Fig. 1f**) with an average diameter of 40 to 150 nm (**Fig. S2a**) based on the statistical analysis of TEM images. Nanoparticle tracking analysis (NTA) showed that the 3T3 exosomes had a narrow size distribution with a mean particle diameter of 116 ± 49 nm (**Fig. S2b**). These exosomes have shown negative charges with a zeta potential of −13.2 ± 2.7 mV measured by dynamic light scattering (DLS). Further characterization by Western blot confirmed that isolated exosomes had specific exosomal markers CD63 and Alix that were scanned versus β-actin (**Fig. S2c**). Overall, these results suggest that exosomes were successfully isolated from 3T3 cell lines and exhibited physical and biological features as nanocarriers for drug delivery.

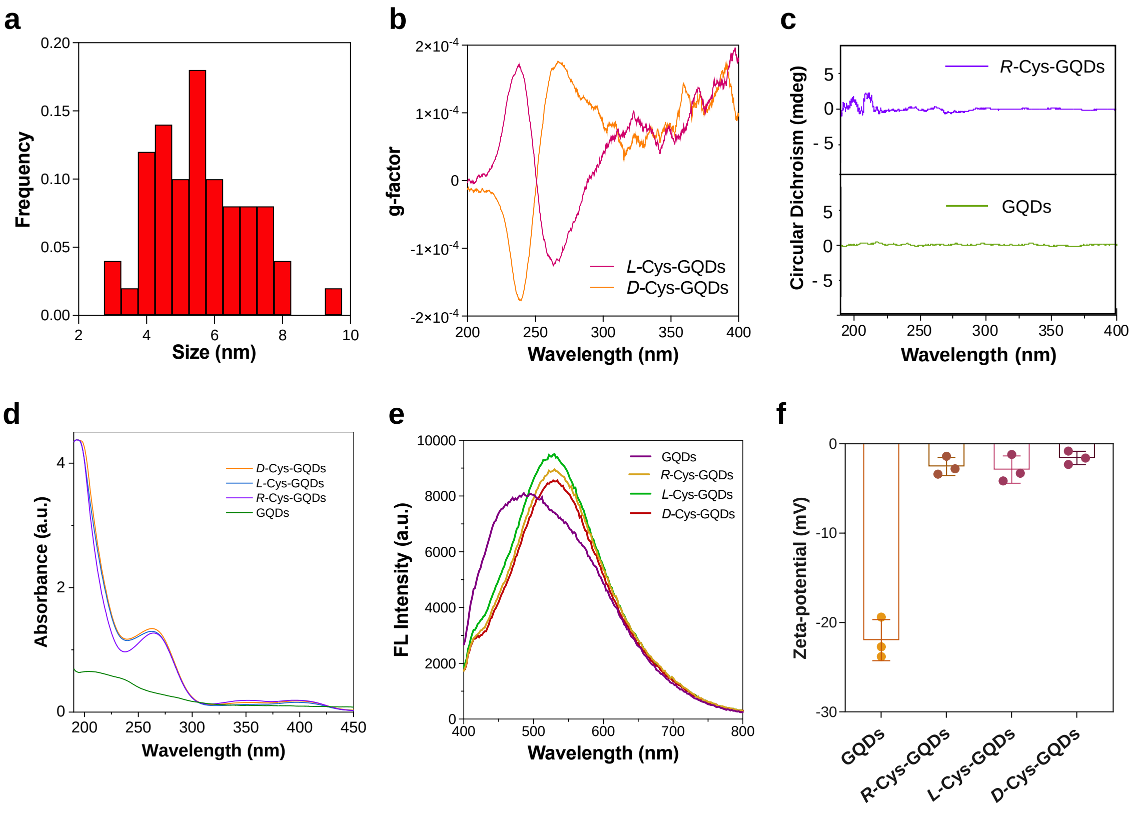


**Figure S1.** Characterizations of GQDs and chiral GQDs. (a) Histograms of size distribution of *L*/*D*-Cys-GQDs, (b) *g*-factor of circular dichroism (CD) spectra for *L*-Cys-GQDs and *D*-Cys-GQDs, (c) CD spectra of GQDs and *R*-Cys-GQDs, (d) UV-Vis spectra and (e) fluorescent spectra of GQDs and *R-*, *L-*, *D*-Cys-GQDs, (f) Zeta-potential of pristine GQDs, *R*-Cys-GQDs and chiral Cys-GQDs.

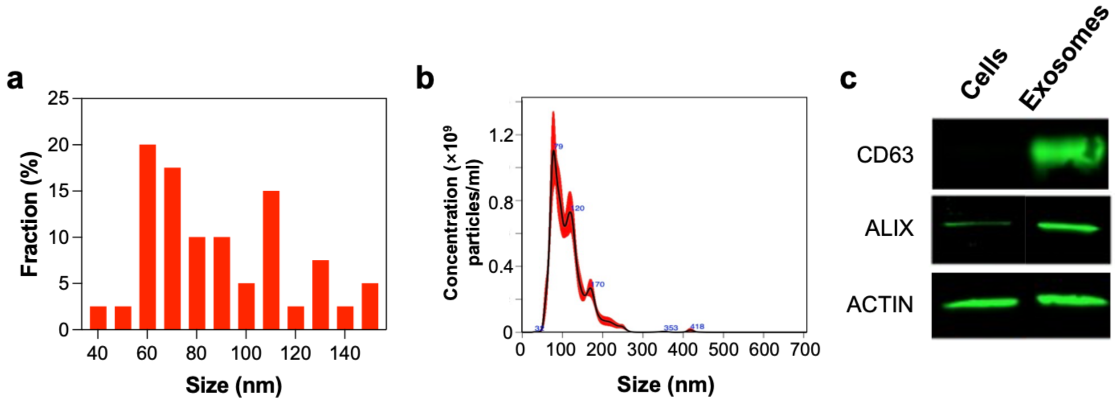


**Figure S2.** Characterization of 3T3 exosomes. a) Size distribution of exosomes based on analysis of TEM images, b) particle number and size distribution of 3T3 exosomes samples were measured with NTA and c) Western blot analyses of exosomal biomarkers (CD63 and Alix).

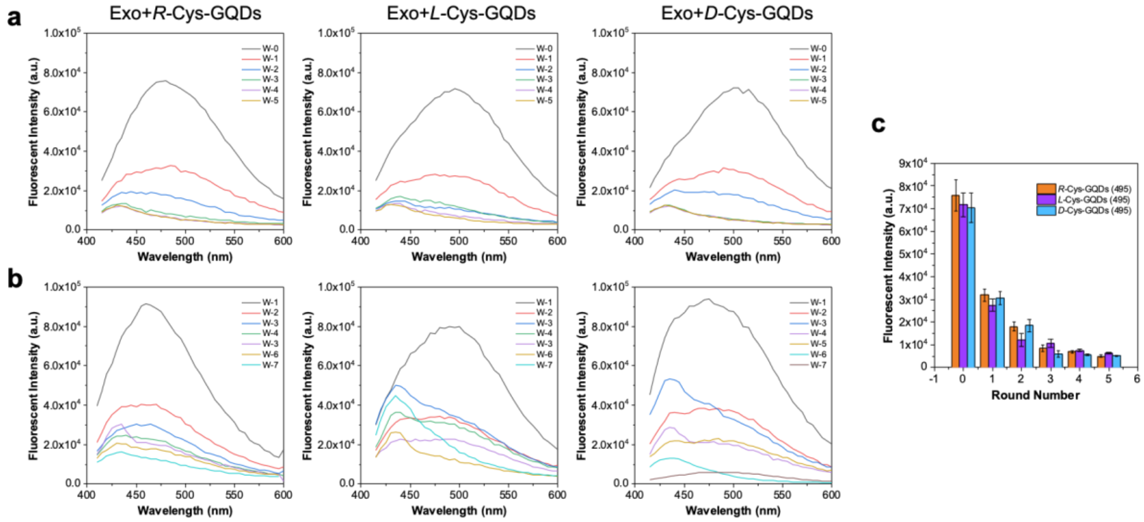


**Figure S3.** Pearmeaction study of chiral Cys-GQDs into exosome. Fluorescence spectra of (a) exosome before and after 7.5 μM *R*-GQDs, *L*-GQDs and *D*-GQDs treatments (Exo: Exosome), and (b) eluent collected with multiple-time (W-1 to W-5, W: Washing times) washes, (c) retention rate.

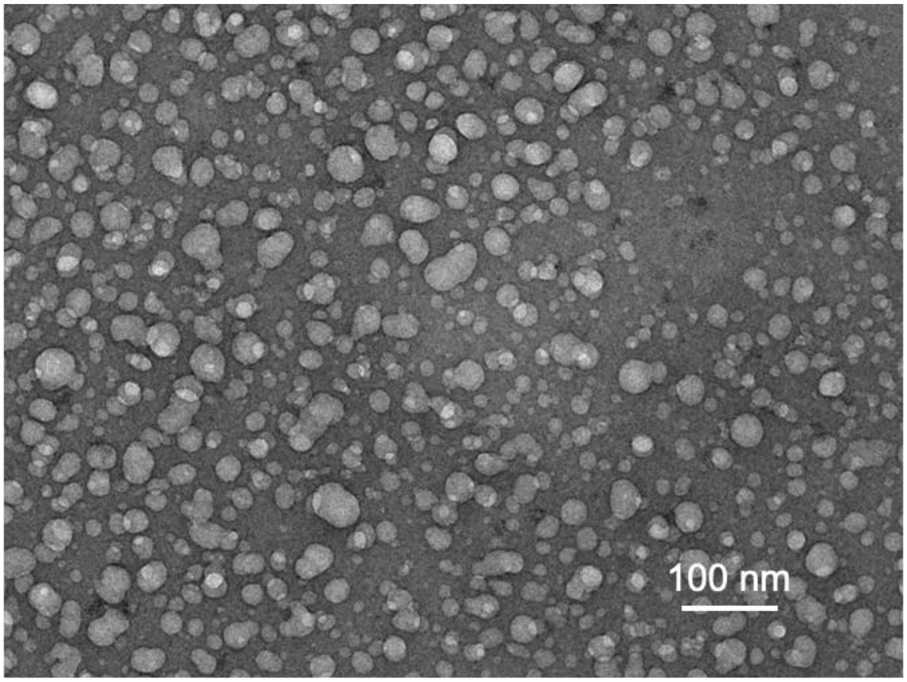


**Figure S4.** TEM image of exosomes treated with *D*-Cys-GQDs.

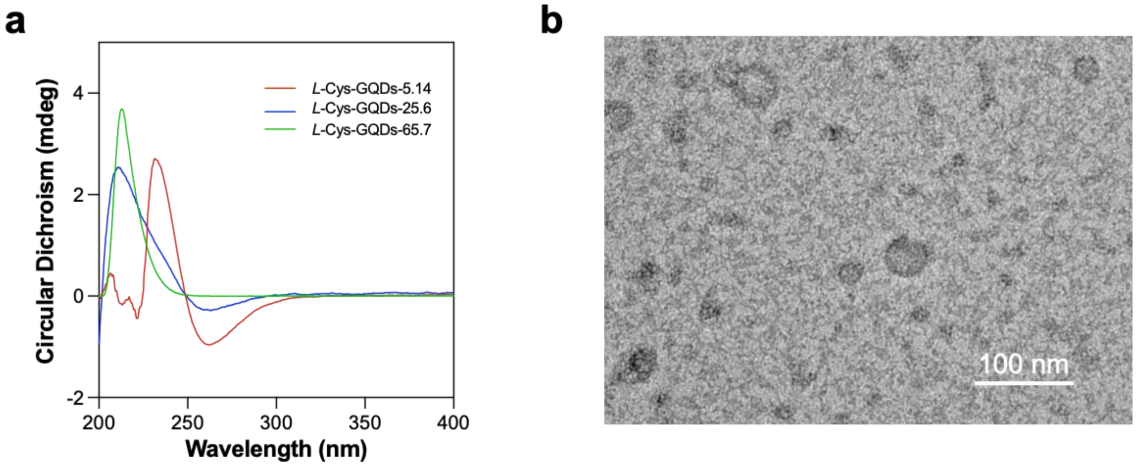


**Figure S5.** Size effect on permeation of chiral GQDs into exosome. (a) CD spectra of *L*-Cys-GQDs with different sizes, and (b) TEM images of damaged exosomes that was treated with *D*-Cys-GQDs with the largest size (15 μM, 65.7 nm).

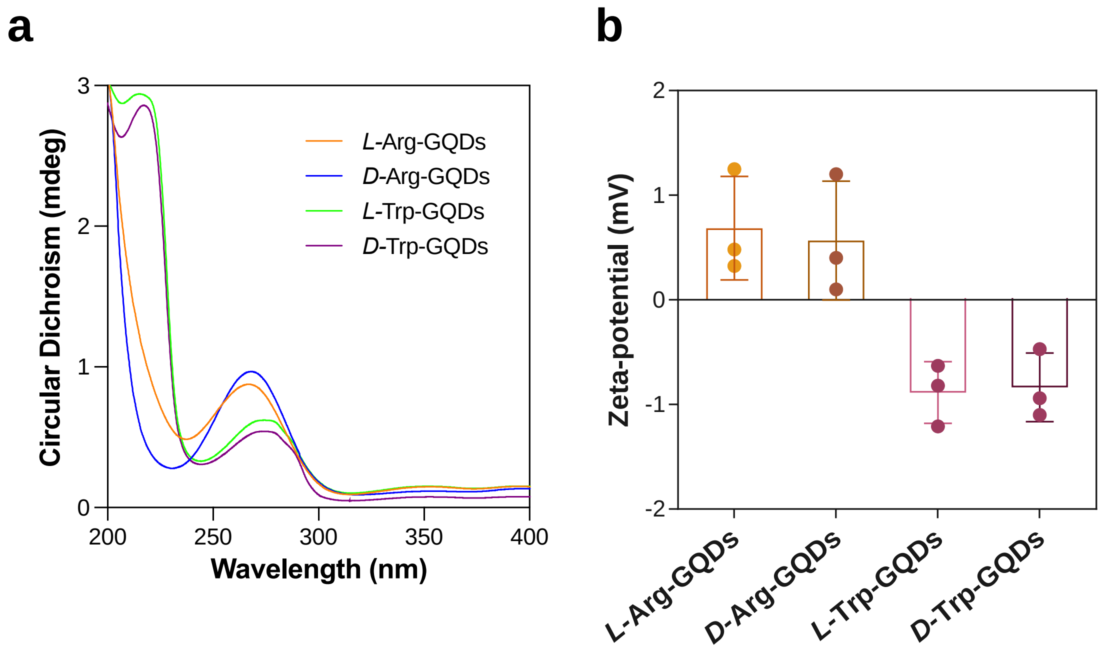


**Figure S6.** (a) UV–vis absorption and (b) zeta-potential (ζ) of the *L/D*-Arg-GQDs and *L/D*-Trp-GQDs.

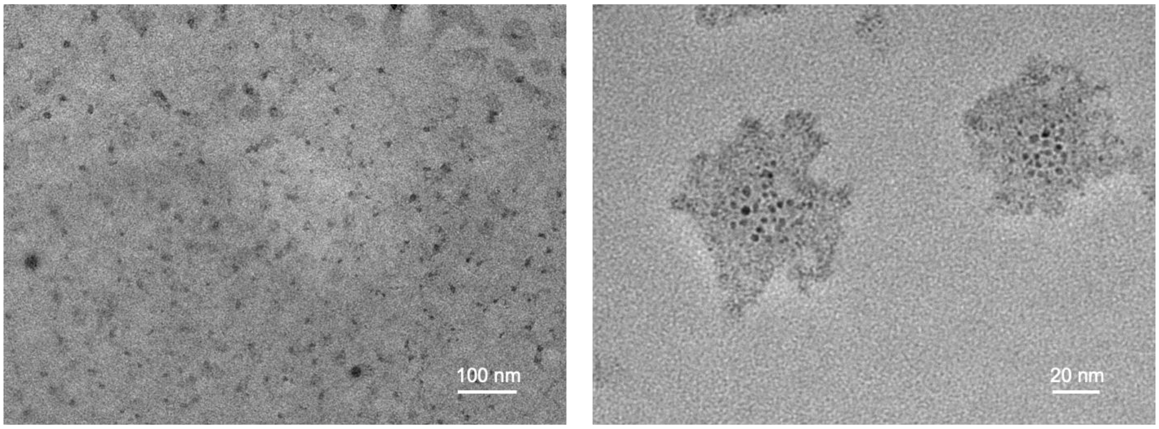


**Figure S7.** TEM images of damaged exosomes that was treated with *D*-Cys-GQDs at the concentration of 30 μM.

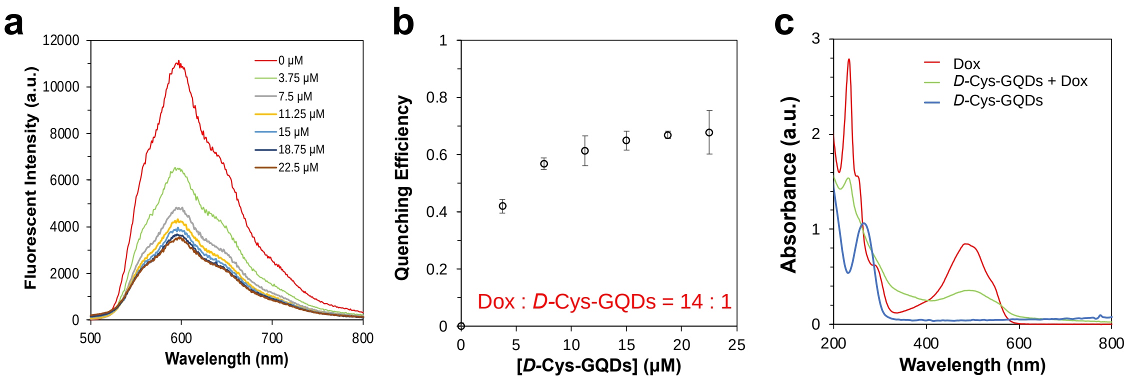


**Figure S8.** The effect of *D*-Cys-GQD concentration on (a) fluorescence intensity and (b) quenching efficiency of Doxorubicin (Dox). (c) UV−vis absorbance spectra of Dox, *D*-Cys-GQDs, and *D*-Cys-GQDs/Dox complex.

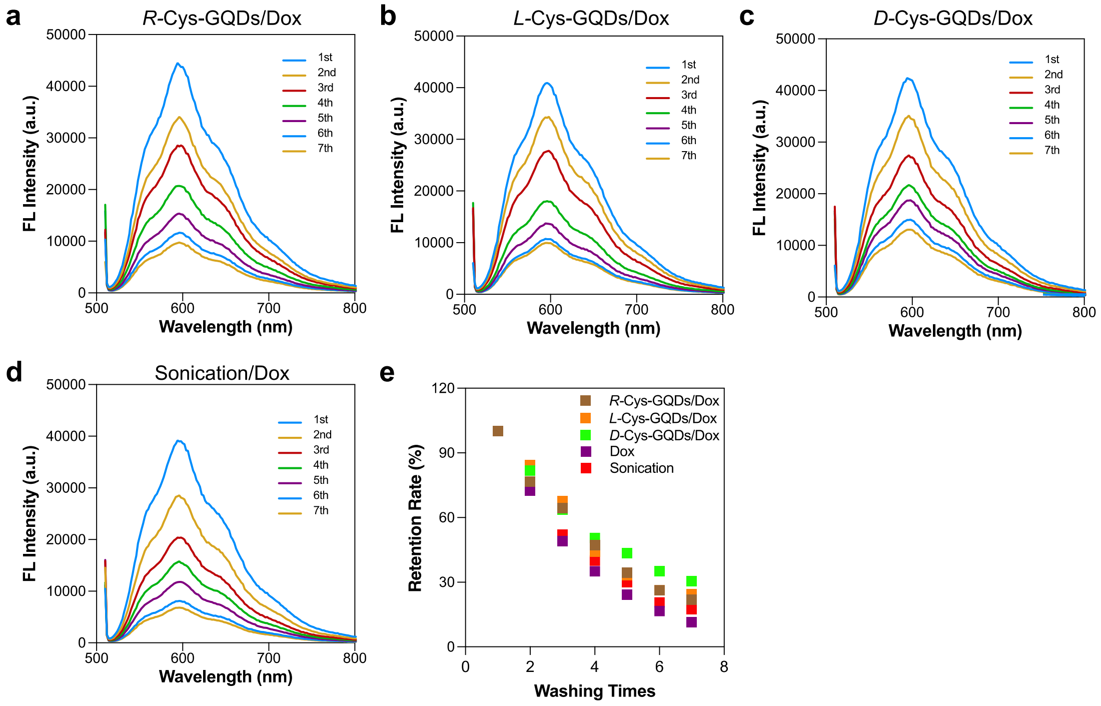


**Figure S9.** Drug loading via chiral Cys-GQDs/Dox into exosome. Fluorescence spectra of exosome after drug loading by (a) *R*-Cys-GQDs/Dox, (b) *L*-Cys-GQDs/Dox, (c) *D*-Cys-GQDs/Dox, (d) sonication treatments and (e) retention rate of Dox in exosomes after washing steps.

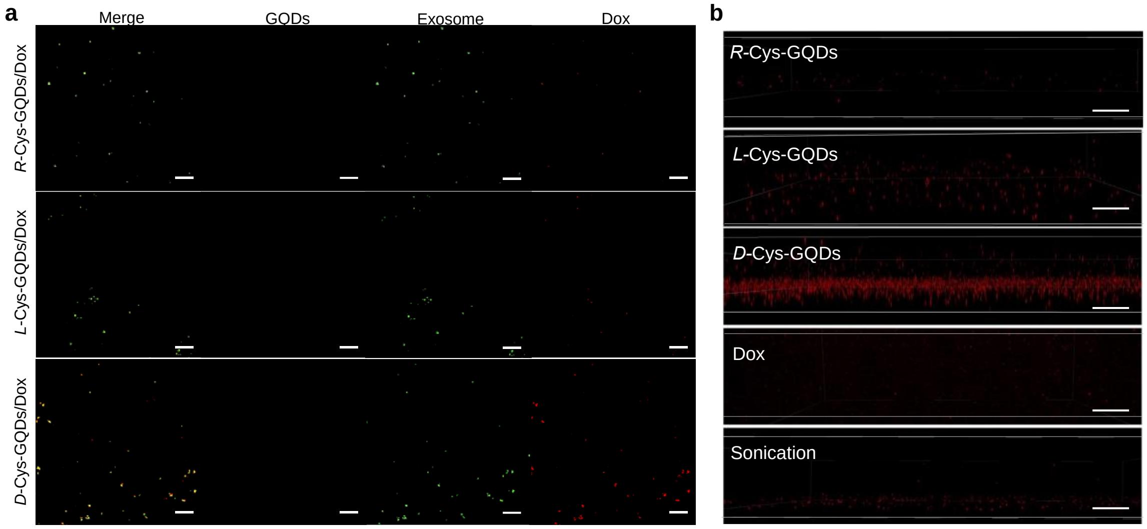


**Figure S10.** Confocal image of free Dox, *R-, L-, D-* Cys-GQDs/Dox and sonication treated exosomes. (a) The confocal image of exosomes loaded by Cys-GQDs/Dox complex (Scale bars: 2 μm) and (b) z-stack confocal images of the Dox (Red) loaded in exosomes (Scale bars: 5 μm).

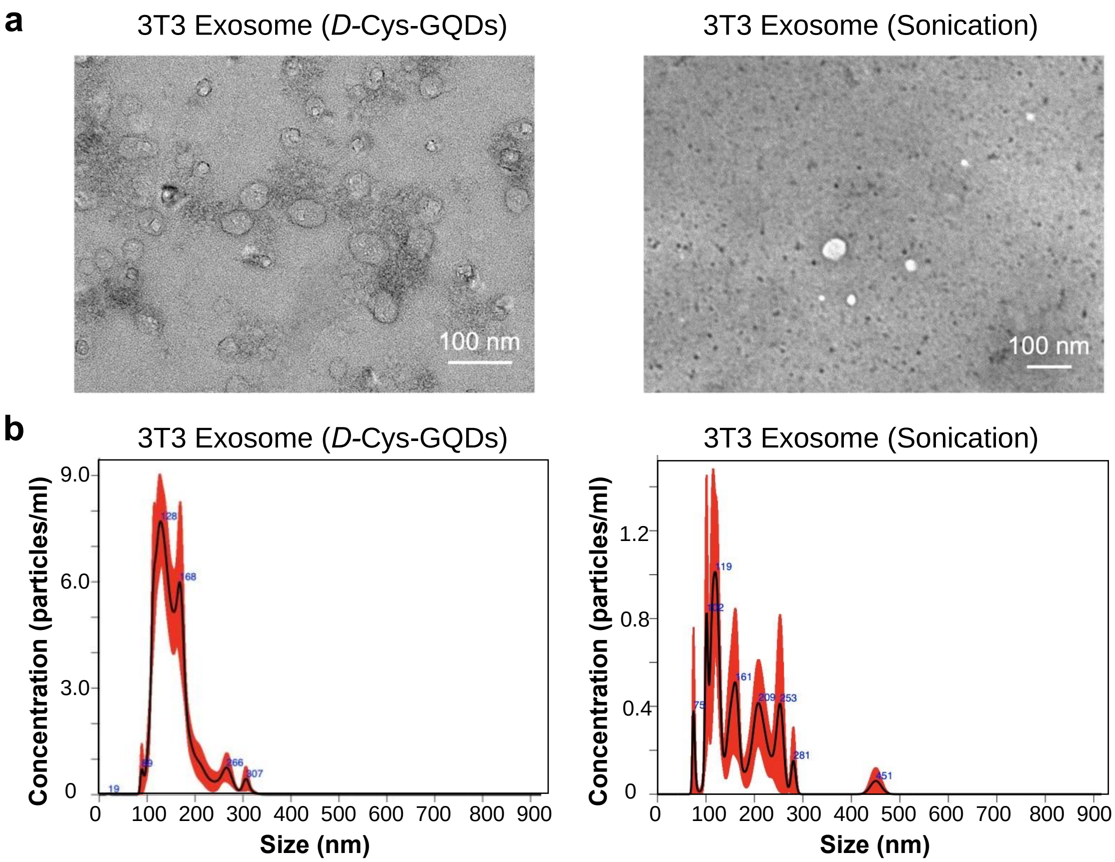


**Figure S11.** Comparison of Dox loading by *D-Cys*-GQDs and sonication. (a) TEM images and (b) particle number and size distribution of 3T3 exosomes samples treated with *D*-Cys-GQDs/Dox and sonication loading method were measured with NTA.

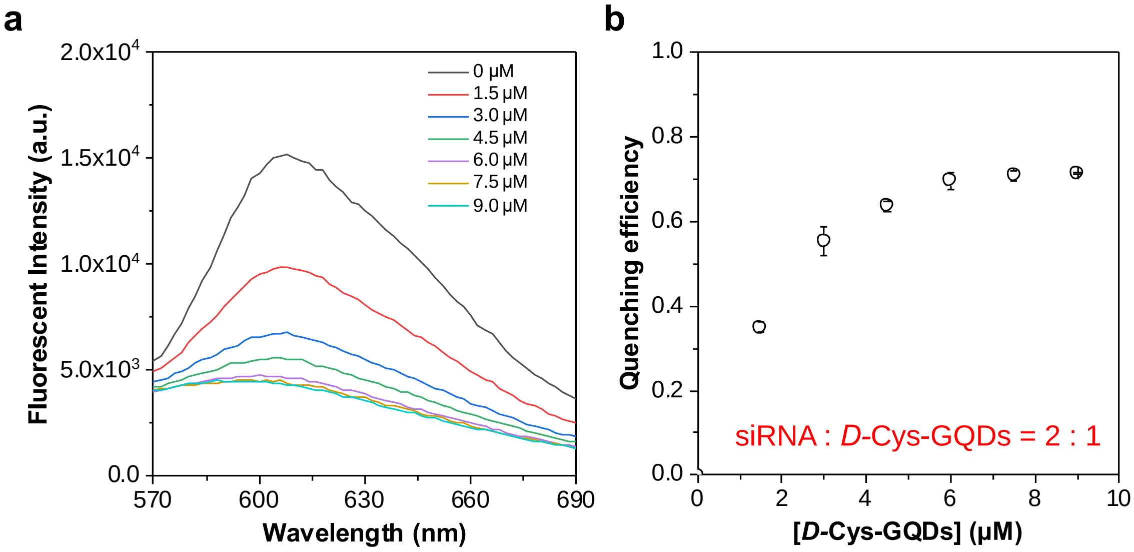


**Figure S12**. The effect of *D*-Cys-GQD concentration on (a) fluorescence intensity and (b) quenching efficiency of 10 μM siRNA.

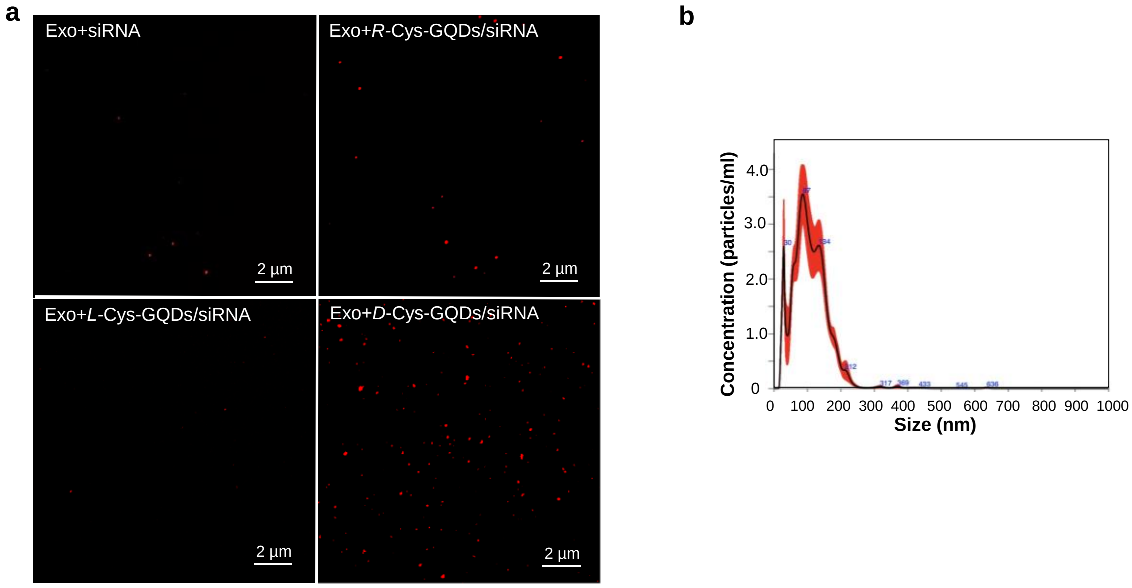


**Figure S13**. The profile of siRNA loaded exosomes. (a) Confocal images of siRNA (Red) loaded exosomes with or without facilitation of *R*-, *L*-, and *D*-Cys-GQDs, and (b) particle number and size distribution of 3T3 exosomes treated with *D*-Cys-GQDs/siRNA was measured with NTA.

### Reference


1. Wang C, Sensale S, Pan Z, Senapati S, Chang H-C. Slowing down DNA translocation through solid-state nanopores by edge-field leakage. *Nature Communications* 2021, **12**(1)**:** 140.
2. Zhang C, Huo X, Zhu Y, Higginbotham JN, Cao Z, Lu X*, et al.* Electrodeposited magnetic nanoporous membrane for high-yield and high-throughput immunocapture of extracellular vesicles and lipoproteins. *Communications Biology* 2022, **5**(1)**:** 1358.
3. Wang C, Senapati S, Chang H-C. Liquid biopsy technologies based on membrane microfluidics: High-yield purification and selective quantification of biomarkers in nanocarriers. *Electrophoresis* 2020, **41**(21-22)**:** 1878-1892.
